# A reservoir of stem-like CD8 T cells in the tumor-draining lymph node maintains the ongoing anti-tumor immune response

**DOI:** 10.1101/2021.01.27.428467

**Authors:** Kelli A. Connolly, Manik Kuchroo, Aarthi Venkat, Achia Khatun, Jiawei Wang, Ivana William, Noah Hornick, Brittany Fitzgerald, Martina Damo, Moujtaba Y. Kasmani, Can Cui, Eric Fagerberg, Isabel Monroy, Amanda Hutchins, Julie F Cheung, Gena G. Foster, Dylan L. Mariuzza, Hongyu Zhao, Weiguo Cui, Smita Krishnaswamy, Nikhil S. Joshi

## Abstract

“Stem-like” TCF1^+^ CD8^+^ T cells (T_SL_) are necessary for long-term maintenance of T cell responses and the efficacy of immunotherapy but, as tumors contain signals that should drive T-cell terminal-differentiation, how these cells are maintained in tumors remains unclear. We found that a small number of TCF1^+^ tumor-specific CD8^+^ T cells were present in tumors throughout development. Yet, most intratumoral T cells differentiated as tumors progressed, corresponding with an immunologic shift in the tumor microenvironment (TME) from “hot” to “cold”. By contrast, most tumor-specific CD8^+^ T cells in tumor-draining lymph nodes (dLNs) had functions and gene expression signatures similar to T_SL_ from chronic LCMV infection and this population was stable over time, despite the changes in the TME. dLN T cells were the precursors of their more-differentiated intratumoral counterparts, and maintenance of TCF1 by intratumoral T cells required continuous migration from dLNs. Finally, T_SL_ CD8 T cells were also present in LNs from lung adenocarcinoma patients, suggesting this population is also relevant in human disease. Thus, we propose that the dLN T_SL_ reservoir has a critical function during tumor development in sustaining antitumor T cells during tumor development and protecting them from the terminal differentiation that occurs in the TME.

## Introduction

Non-small lung cancer (NSCLC) is the deadliest cancer (*2*), but immune checkpoint inhibitors (ICIs), like anti-PD-1 and anti-PD-L1, have provided durable responses in ∼20% of treated NSCLC patients (*3*). Several parameters correlate positively with response, including the presence of an immunologically “hot” tumor microenvironment (TME; contains infiltrating CD3+ T cells; also called T-cell inflamed), infiltration of PD-1^+^ CD8^+^ T cells, PD-L1 immunostaining on tumor and immune cells, and increased tumor mutational burden/neoantigens (*4–7*). These observations are in line with the idea that PD-1 blockade potentiates the function of “exhausted” PD-1^+^ tumor-infiltrating CD8^+^ T cells in hot tumors (*8*). By contrast, patients with immunologically “cold” tumors (also called “immune deserts” or “non-T-cell inflamed”) respond poorly to immunotherapy, and the status of their anti-tumor CD8 T cell response is uncertain (*4–7*). As response rates to immunotherapy are low, particularly for patients with cold tumors, a better understanding of the CD8 T cell immunobiology associated with hot and cold tumors is paramount.

CD8 T cell exhaustion is a progressive process of terminal differentiation (*9–12*). Exhausted T cells (T_EX_) can be characterized by the loss of proliferative potential and effector functions (ability to produce TNFα and IFNγ), as well as increased expression of several inhibitory receptors (*e.g.,* PD-1 and Tim3) and transcription factors (*e.g.,* Blimp-1, Tox, and Eomes) (*9, 12–30*). T_EX_ cells are derived from less differentiated precursors, including PD-1^mid^ CXCR5^+^ SLAMF6^+^ TCF1^+^ “stem-like” T (T_SL_) cells (*12, 21, 24, 31-40*). TCF1^+^ T_SL_ cells have at least two functions in chronic immune responses: maintaining the ongoing T cell response and mediating therapeutic responses to PD-1 blockade (*24, 31, 33, 34, 41*). Expression of TCF1 is necessary for both functions (*12, 34*), and thus, provides an important tool for identifying T_SL_ cells.

TCF1^+^ CD8 T cells are present in tumors, and their presence correlates with better outcomes following immunotherapy (*10, 12, 21, 24, 31, 33, 34, 37, 40-43*). Yet, tumors are rich in signals that promote the exhaustion of CD8 T cells, like persistent antigen exposure (*15, 44*). This raises a fundamental question: how are intratumoral TCF1^+^ CD8 T cells maintained over the months-to-years of natural tumor development? Moreover, because TCF1 expression is required for maintenance of T cell populations, do immunologically cold tumors result from a loss of intratumoral TCF1^+^ CD8 T cells.

The KP (Kras^lox-stop-lox (lsl)-G12D/+^;p53^flox/flox^) model is a genetically engineered mouse (GEM) model of cancer that faithfully recapitulates the histological, transcriptomic, epigenomic, and genetic features of a developing human lung adenocarcinoma and has played a fundamental role in our understanding of how human lung cancer develops (*45–48*). Tumors in the KP model can be programmed to express neoantigens, which allows for the investigation of tumor-specific T cell responses in developing tumors (*49–59*). Early neoantigen-expressing KP lung tumors are infiltrated by tumor-specific CD8 T cells and have an immunologically hot TME. However, as tumors develop, they take on an immunologically cold TME, with T cells being excluded from the tumor parenchyma and restricted to tertiary lymphoid structures (TLSs) (*49, 60*). Thus, the KP model provided us with an opportunity to investigate the differences between T cells from hot and cold tumor microenvironments, and to assess the mechanisms for how tumor-specific CD8 T cells are maintained over the course of tumor growth.

## Results

### Tumor-specific TCF1^+^CD8^+^ T cells are present throughout disease progression

To study tumor-specific CD8 T cell responses in neoantigen-expressing lung tumors, we infected KP-NINJA mice (KP x *R26-NINJA* x *CCSP-rtTA Tg*) intratracheally (i.t.) with adeno- or lentiviruses expressing Cre alone (Ad5mSPC-Cre or lenti-Cre) (**Figure 1A**)(*59, 61*). In this model, Cre-expression in infected lung epithelial cells activates *Kras G12D* and eliminates *Trp53*. Expression of the neoantigens in tumor cells (GP33-43 and 66-77 from the LCMV glycoprotein) is initiated by doxycycline and tamoxifen (Dox/Tam) treatment starting 7-10 days post infection (**Figure 1B**). Tumors develop progressively after initiation with tumor-specific CD8 T cells consistently detectable as early as 8 wks pi and large, macroscopically visible lung tumors by 16-20 wks pi (data not shown).

**Figure 1:**
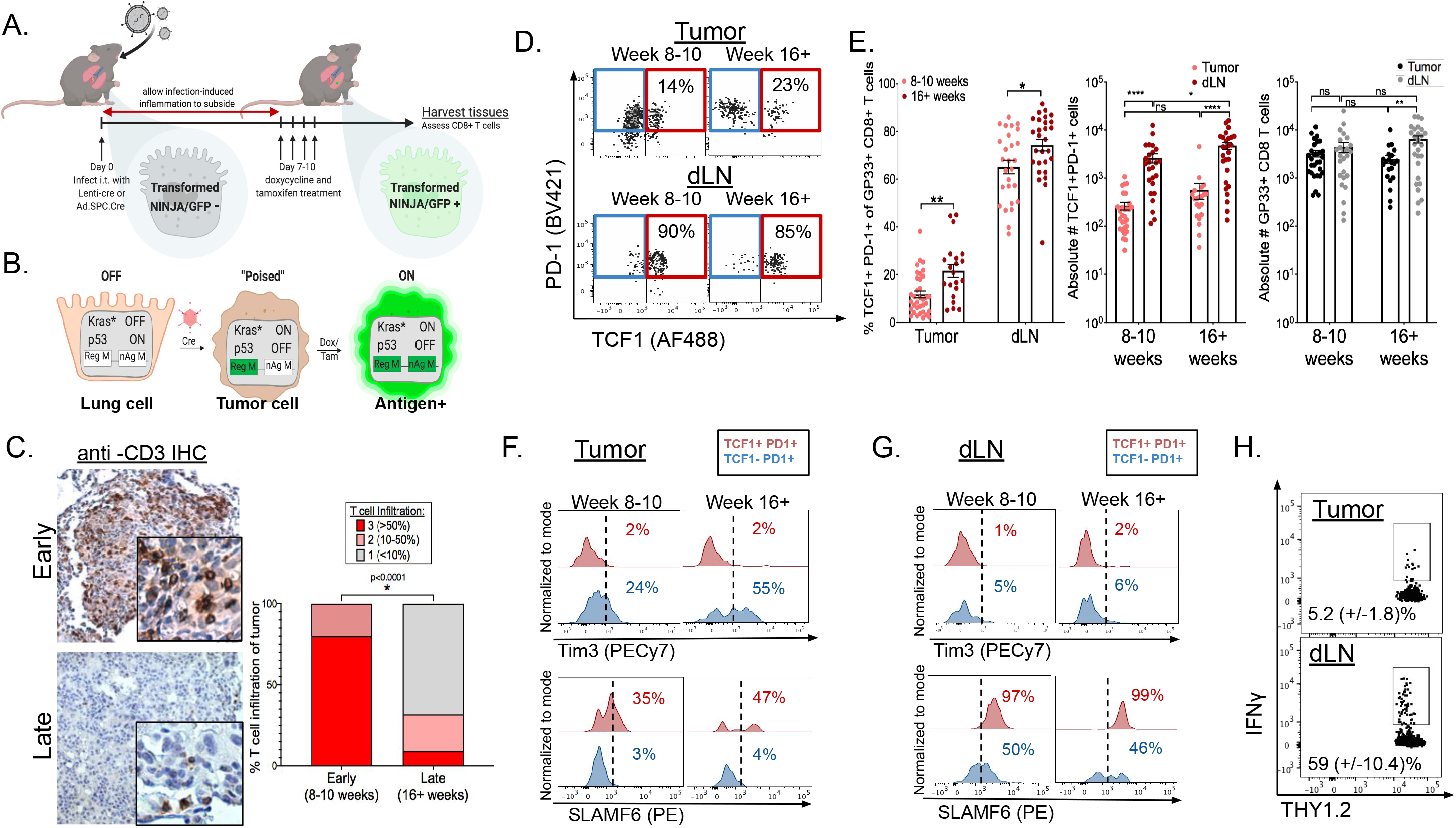
Tumor-specific TCF1^+^CD8^+^ T cells are present throughout disease progression in autochthonous KP-NINJA lung tumors. (A) Experimental setup of KP-NINJA tumor induction in which doxycycline and tamoxifen treatment is administered on days 7-10 after initial infection. (B) Schematic detailing the genetic recombination events in KP-NINJA system. In KP-NINJA mice, i.t. administration of adeno- or lentiviruses expressing Cre alone (Ad5mSPC-Cre or lenti-Cre), activates Kras G12D and eliminates p53 in lung epithelial cells. In cancer cells, but not other cells, Cre also causes a permanent DNA inversion in the FLPoER-encoding regulatory module (Reg M) of the NINJA allele to “poise” Reg M (*59*). When cells have the Reg M in the poised configuration, subsequent Dox/Tam treatment drives FLPoER activity, causing a permanent inversion in the neoantigen module (nAg M) and allowing the expression of neoantigens encoding GFP (contains neoantigens from the lymphocytic choriomeningitis virus (LCMV) glycoprotein: GP33-43 and 66-77). Here, Cre-exposed cells remain neoantigen negative until Dox/Tam administration, allowing for temporal dissociation of tumor initiation and neoantigen induction. (C) De-identified, anti-CD3 immunohistochemistry stained KP-NINJA tumor-bearing lungs were scored blindly for level of T cell infiltration (3=>50%;2=10-50%;1=<10%). Scores were compared between early (8-10 weeks post infection; n=10) and late (16+ weeks post infection; n=22) tumors. p=<0.0001 by unpaired t-test. (D) Representative flow cytometry dot plots displaying extracellular expression of PD-1 and intracellular expression of TCF1 on tissue GP33-specific CD8^+^ T cells from the tumors (top) and dLNs (bottom) at early (8-10 weeks) and late (16+ weeks) time points after infection, cells pre-gated as in (Figure S1A). (E) Left panel: Percent TCF1^+^PD-1^+^CD8^+^ T cells of total GP33-specific CD8^+^ T cells (**p=0.0008 tumor; *p=0.0160 dLN). Center panel: Absolute numbers of TCF1^+^PD-1^+^CD8^+^ T cells (*p=0.0415; ****p=<0.0001), and absolute number of total GP33-specific (GP33 loaded MHC I tetramer^+^) CD8^+^ T cells (**p=0.0041) in tumors (black) and dLNs (gray). (F-G) Representative histograms displaying extracellular Tim3 expression (top) and SLAMF6 expression (bottom) of TCF1^-^PD-1^+^ vs. TCF1^+^PD-1^+^ tumor-specific CD8^+^ T cells at early (8-10 weeks) and late (16+ weeks) after infection in tumors (F) and dLNs (G). Data from 8 (early) and 5 (late) independent experiments: n=29 tumors at 8-10 weeks p.i.;n=21 tumors at 16+ weeks p.i.; n=25 dLN at 8-10 weeks p.i.;n=27 dLN at 16+ weeks p.i. Statistics based on two-tailed, unpaired t-test. Mean and SEM reported in text. (H) Representative dot plots showing *ex vivo* IFNγ production of tumor (top) and dLN (bottom) cells following GP33-41 peptide restimulation. Representative of 2 independent experiments; n=7 **p=0.005. Note: similar results to 1D were seen when tumors were programmed to express neoantigens with lentiviruses containing the neoantigens in NINJA, demonstrating that T cell differentiation was independent of the method used for neoantigen programing (**Figure S1D**).

Previous studies have shown a shift from hot to cold tumors between early and late tumors (*49, 60*). To confirm this, we quantified T-cell infiltration by CD3 immunohistochemical (IHC) staining in 8 and 16-20 wk tumors. Tumors in the KP-NINJA model underwent a transition from an immunologically hot to an immunologically cold TME (**Figure 1C**). This shift was not due to loss of tumor antigens, as cell lines made from 20 wk tumors expressed neoantigens had inducible expression of MHC class I and PD-L1 and elicited GP33- and GP66-specific CD8 and CD4 T cell responses (respectively) after transplant (data not shown). Thus, we hypothesized that the shift from the hot to cold TME might reflect a global exhaustion of tumor-specific CD8 T cells in animals at late timepoints.

To test our hypothesis, we analyzed the expression of TCF1 and PD-1 on lung-tissue tumor-specific CD8^+^ T cells from early (8-10 week) and late (16+ week) tumor-bearing KP-NINJA mice (**Figure 1D**; tumor-specific CD8 T cells identified with GP33-loaded MHC tetramer^+^ cells; gated based on FMO controls for each sample as in **Figure S1B**). Surprisingly, at both time points studied, there was an appreciable population of TCF1^+^PD-1^+^ CD8 T cells (mean of 12(+/−1.4)% and 22(+/−2.6)%, respectively; **Figure 1E left**). Moreover, their number did not change substantially as tumors progressed (**Figure 1E center;** gates based on PD-1 and TCF1 staining in GP33^-^ population of i.v.CD45^-^ THY1.2^+^ CD8^+^ T cells and TCF1 FMO controls, **Figure S1C**). Yet, despite this, there was a ∼2x increase in the fraction of terminally differentiated Tim3^+^ TCF1^-^ PD-1^+^ CD8 T cells between early and late tumors (**Figure 1F top**). Less than 50% of the TCF1^+^ PD-1^+^ CD8 T cells in tumors also expressed the T_SL_ marker SLAMF6 at either early or late time points (**Figure 1F bottom**). Together, these data suggested that the cold tumor phenotype in advanced KP tumors might not be due to the loss of TCF1^+^ CD8 T cells.

The presence of tumor-specific TCF1^+^PD-1^+^ CD8 T cells throughout tumor development was intriguing because it suggested that mechanisms exist for the maintenance of this population in tumors. We reasoned this could be due to local maintenance of TCF1^+^ cells and/or migration from a distal site. Thus, we assessed tumor-dLNs from KP-NINJA mice for TCF1^+^PD-1^+^ CD8 T cells. Critically, ∼65(+/−2.8)% and ∼74(+/−2.4)% of the tumor-specific CD8^+^ T cells in the dLN were TCF1^+^ and PD-1^+^ at 8-10 and 16+ weeks post tumor initiation, respectively (**Figure 1D-E**). Numerically, this amounted to ∼10-fold more TCF1^+^PD-1^+^ GP33-specific CD8 T cells in dLNs compared to tumors (mean number of T_SL_= 264+/−48 in tumor vs. 2,630+/−573 in dLN 8-10 weeks p.i. and 567+/−204 in tumor vs. 4,730+/−807 in dLN at 16+ weeks p.i.) (**Figure 1E center**). This population in the dLN appeared phenotypically stable over time and even remained Tim3^-^ throughout tumor development (**Figure 1F top**). Furthermore, and in contrast to the TCF1^+^PD-1^+^ CD8 T cells in tumors, the TCF1^+^PD-1^+^ CD8 T cells in early and late dLNs were SLAMF6^+^ (**Figure 1F bottom**). Functionally, while a portion of T cells from tumors produced IFNγ after GP33-41 peptide re-stimulation *ex vivo* (5.2 +/− 1.8%; normalized to the frequency of GP33-loaded Tetramer^+^ CD8 T cells in lung tissue), the T cells from dLN appeared to have an enhanced functional capacity, as a much higher proportion were IFNγ^+^ after re-stimulation (59 +/−10.4%; normalized to the frequency of GP33-loaded Tetramer^+^ CD8 T cells in dLN) (**Figure 1G**). Together these data demonstrated that most tumor-specific CD8 T cells in the dLN were TCF1^+^ and SLAMF6^+^, and that a significant fraction of these cells had functional capacity.

### TCF1^+^CD8^+^ T cells are enriched in dLNs in an orthotopic lung tumor model

To validate our results in a second tumor model, we established an orthotopic lung tumor transplant model with a cell line (KPN1) derived from an advanced tumor in a KP-NINJA mouse. This tumor line grew well in immunodeficient mice and contained very few chromosomal copy number alterations compared to longer established KP cell lines (data not shown). Moreover, this line elicited responses from endogenous GP33-specific CD8 T cells after orthotopic transplant. Analysis of GP33-specific CD8 T cells in dLN and tumors at days 10 and 20 post-transplant (p.t.) showed that TCF1^+^PD-1^+^ T cells were present in the lungs at both time points, but that there was a significant population of TCF1^+^PD-1^+^ T cells in the dLN (**Figure S2**). Moreover, their frequency and number were greater in the dLN. Thus, using a second model, we confirmed that the dLN was a site with significant enrichment of tumor-specific TCF1^+^ PD-1^+^ CD8^+^ T_SL_ cells.

### Comparing the differentiation of tumor-specific CD8 T cells to CD8 T cells from chronic LCMV infection

To better understand the differentiation of tumor-specific CD8 T cells at early and late timepoints, we sorted endogenous GP33-specific CD8 T cells from the dLNs and lung tissues (i.v.CD45-) of KP-NINJA mice 8 (early) and 17 weeks p.i. (late) and performed single cell RNA sequencing (RNAseq) with paired V(D)J sequencing of the T cell receptor (TCRseq; **Figure 2A**). The lineage of exhausted CD8 T cells has been well described in the context of chronic LCMV infection and a strength of the KP-NINJA model is that endogenous GP33-specific CD8 T cells recognize the same antigenic peptides as those present in acute (Armstrong) and chronic (Clone 13) LCMV. We sorted endogenous GP33-specific CD8 T cells from spleens of C57BL/6 mice 28 days after infection with LCMV Clone 13 or LCMV Armstrong and performed single cell RNAseq and TCRseq. We then directly compared the transcriptomes of the sorted anti-tumor and anti-viral T cells.

**Figure 2:**
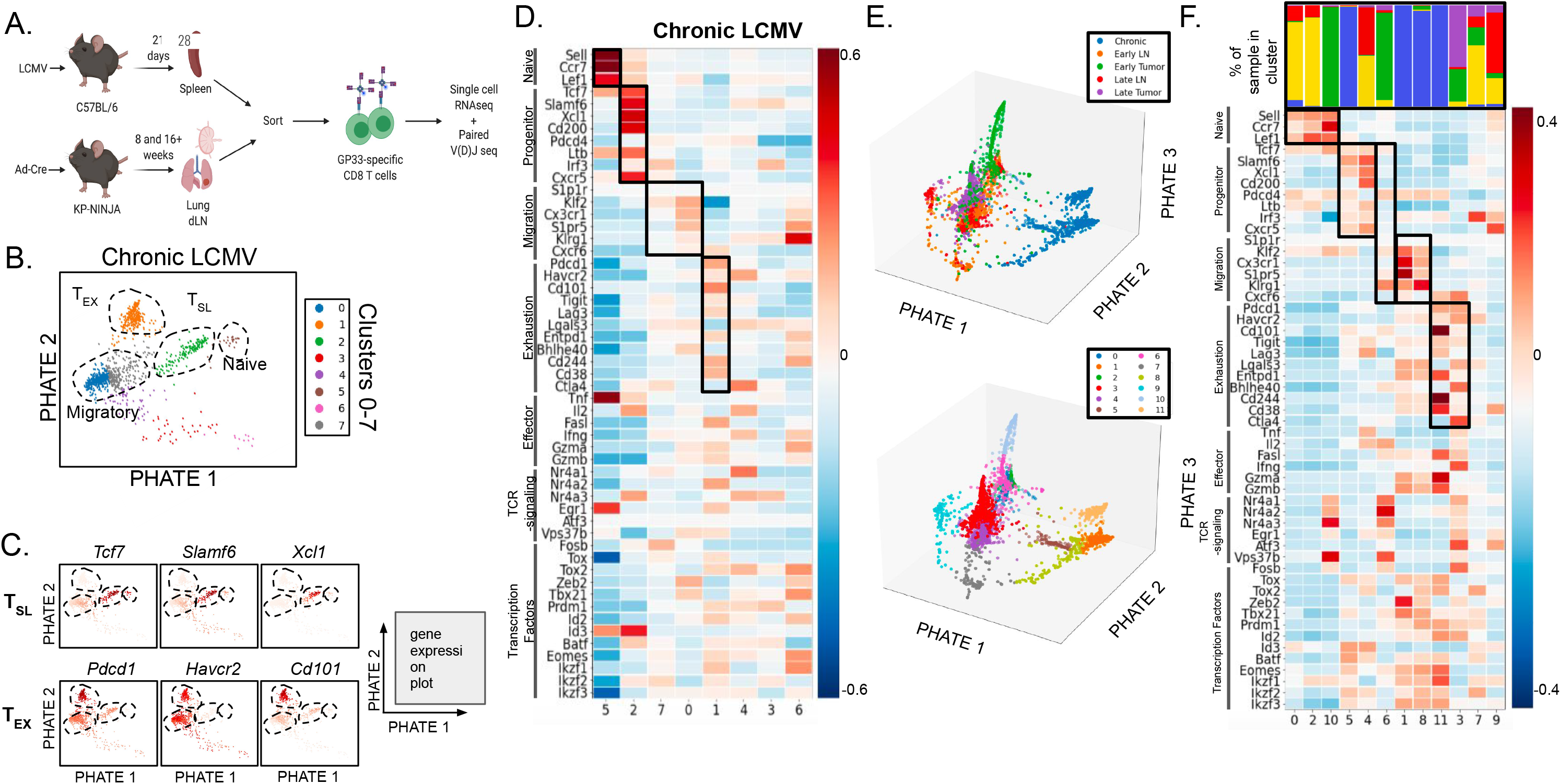
Tumor-specific TCF1^+^PD1^+^ CD8 T cells in tumor and dLN resemble T_SL_ cells in chronic LCMV. (A) Experimental protocol schematic. Spleens were harvested from C57BL/6 mice 28 days following infection with LCMV-Clone 13 (or LCMV-Armstrong – Figure S3H-J). Likewise, tumors and dLNs were harvested from tumor-bearing KP-NINJA mice 8 and 17 weeks after intratracheal infection with Ad5mSPC-Cre and doxycycline and tamoxifen treatment. GP33-specific CD8^+^ T cells were sorted from tissues (i.v.CD45^-^ CD8^+^GP33-loaded MHC I tetramer^+^) and submitted to the Yale Center for Genome Analysis for single-cell RNA and TCR sequencing. Single cell RNA-sequencing data was demultiplexed using Cell Ranger 3.0 Software and then further analyzed using Python. (B) Single-cell RNAseq data from GP33-specific CD8^+^ T cells from chronic LCMV Clone 13 infection were analyzed. 1,185 cells analyzed and 11,595 genes detected. Cell clusters for GP33-specific CD8^+^ T cells from chronic LCMV Clone 13 infection were visualized using PHATE maps and colored based on clustering into arbitrary 7 clusters. (C) Naïve-, T_SL_-, and T_EX_-like populations (dotted lines) were identified based on the expression profiles of key genes associated with naïve (*Sell, Lef1, Ccr7*), stem-like precursor (*Tcf7, Slamf6, Xcl1*), and terminally exhausted (*Pdcd1, Havcr2, Cd101*) CD8^+^ T cells. See Figure S3A for analysis of additional genes in the following categories: progenitor-related, migration-related, and exhaustion-related, effector molecules, transcription factors, TCR signaling, and naïve-related. (D) Heat map based on gene expression of naïve related (*Sell, Ccr7, Lef1*), progenitor-related (*Tcf7, SLAMF6, Xcl1, Cd200, Pdcd4, Ltb, Irf3, Cxcr5*), migration-related genes (*Cxcr6, Klf2, Cx3cr1, S1pr5)*, exhaustion-related (Pdcd1, Havcr2, Cd101, Tigit, Lag3, Lgals3, Entpd1, Bhlhe40, Cd244, Cd38, Ctla4), as well as key effector cell-associated genes (*Klrg1, Tnf, Il2, Fasl, Ifng, Gzma, Gzmb*), genes associated with intracellular TCR signaling (*Nr4a1, Nr4a2, Nr4a3, Egr1, Vps37b, Fosb*), and key transcription factors (*Fosb*, *Tox*, *Tox2*, *Zeb2*, *Tbx21*, *Prdm1*, *Id2*, *Id3*, *Batf*, *Eomes*, *Ikzf1*, *Ikzf2*, *Ikzf3*). (E) 3-dimensional PHATE map of GP33-specific CD8 T cells from chronic LCMV infection, early dLN, late dLN, early tumor, and late tumor were combined and divided into 12 (0–11) clusters (refer to Figure S3C). (F) Heat map from combined analysis with distribution of samples for each clusters shown above. Gene expression of naïve related (*Sell, Ccr7, Lef1*), progenitor-related (*Tcf7, SLAMF6, Xcl1, Cd200, Pdcd4, Ltb, Irf3, Cxcr5*), migration-related genes (*Cxcr6, Klf2, Cx3cr1, S1pr5)*, exhaustion-related (Pdcd1, Havcr2, Cd101, Tigit, Lag3, Lgals3, Entpd1, Bhlhe40, Cd244, Cd38, Ctla4), as well as key effector cell-associated genes (*Klrg1, Tnf, Il2, Fasl, Ifng, Gzma, Gzmb*), genes associated with intracellular TCR signaling (*Nr4a1, Nr4a2, Nr4a3, Egr1, Vps37b, Fosb*), and key transcription factors (*Fosb*, *Tox*, *Tox2*, *Zeb2*, *Tbx21*, *Prdm1*, *Id2*, *Id3*, *Batf*, *Eomes*, *Ikzf1*, *Ikzf2*, *Ikzf3*) are included. For early dLN, 1,742 cells were analyzed and 12,116 genes were detected. For late dLN, 876 cells were analyzed and 11,595 genes were detected. For early tumor, 806 cells were analyzed and 11,749 genes were detected. For late tumor, 731 cells were analyzed and 11,150 genes were detected. Single cell RNA-sequencing data is representative of n=3 pooled samples.

First, we analyzed our single cell RNAseq data from chronic LCMV infection to identify clusters that fit the previous descriptions for naïve, “precursor-exhausted” T_SL_, “transitory”/migratory effector T cells, and terminally exhausted T_EX_ cells (*10-12, 21, 24, 31, 34, 35, 40, 43, 62, 63*). We leveraged PHATE embedding to visualize our analyses. PHATE is an unbiased clustering tool optimized to learn and represent transitions between cell states from high-dimensional single-cell data sets without imposing strong assumptions on the structure of the data (*64*). Similar to UMAP projections, the organization of clusters and the relative distances between clusters on PHATE embeddings have meaning (*i.e.,* closely related clusters are located in closer physical proximity). Unbiased clustering analysis on the GP33-specific CD8 T cells from chronic infection revealed 8 clusters (**Figure 2B**). We analyzed the PHATE embedding for the expression of select genes that have previously been associated with T cell subsets in chronic LCMV (naïve related, progenitor-related, migration-related, and exhaustion-related). Genes encoding T-cell effector molecules (to assess function), transcription factors associated with CD8 T cell differentiation, and genes associated with TCR-signaling were also included (**Figure 2C and S3A**) (*9, 11, 12, 26-29, 31, 33, 35, 39, 43, 65-71*). Based on patterns of gene expression, we identified clusters that best represented naïve (5), T_SL_ (2), migratory (7, 0), and T_EX_ (1) cell populations (**Figure 2B and D**). Moreover, beyond expression of the selected genes, unbiased PHATE embedding recapitulated the expected lineage relationships between the clusters, suggesting progressive differentiation from T_SL_ to migratory and then T_EX_ cells.

We next turned our focus to analysis of the individual samples from tumors and dLNs (**Figure S3B-E**). Unbiased cluster analysis of the individual samples suggested that each contained several populations, including a cluster cells with a strong naïve T cell signature, which we suspected were contaminating CD8 T cells in our GP33-specific T cell sorts. Using TCRseq data, we confirmed that the naïve T cell clusters were comprised entirely of T cells with unique TCR sequences (*i.e.,* singlets), while the non-naïve T cell clusters contained T cells with TCRs that were “shared” with other T cells in the sample (**Figure S3F**). Analysis of the dLN T cells suggested relatively uniform expression of progenitor-, migration-, and exhaustion-related genes and genes for effector molecules and transcription factors across most cells (**Figure S3B and D**). By contrast there was more heterogeneity in expression of these genes amongst T cells in the tumor samples.

To compare T_SL_ and T_EX_ T cell subsets between the tumors, LNs, and chronic LCMV samples, we co-embedded all five data sets (early and late dLNs and tumors and chronic infection) and visualized their co-embedding by PHATE (**Figure 2E and S3G**). This analysis identified 12 clusters, and based on their expression the signature panel genes, we identified clusters of naïve (0, 2, 10), T_SL_ (5, 4, 6), migratory (8, 1) and T_EX_ (11, 3) cells (**Figures 2F**).

The 3 T_SL_ clusters included one that was predominantly made up of cells from chronic infection (5) and one that predominantly had cells from the two dLN samples (4). These clusters had similar gene expression patterns, although there were notable differences in *Cd200* and *Pdcd4* (**Figure 2F**). Interestingly, Cluster 6 was predominantly made up of cells from early tumors and had low expression of most of progenitor-related signature genes, with the exception of *Tcf7*. These data suggested that TCF1^+^ SLAMF6^+^ tumor-specific CD8 T cells in dLNs were closer in gene expression to T_SL_ cells from chronic infection than their TCF1^+^ counterparts in tumors. Notably, the placement of these clusters in a 3-dimensional PHATE embedding confirmed the close proximity between T_SL_ from chronic infection (cluster 5, brown) and T_SL_ in dLNs (cluster 4, purple) (**Figure 2E**).

The 2 T_EX_ clusters were predominantly made up of cells from chronic infection (11) and early/late tumors (3) (**Figure 2E**). Clusters 11 and 3 shared high expression of most, but not all, of the exhaustion signature genes, suggesting that the TCF1^-^ PD-1^hi^ Tim3^+^ tumor-specific CD8 T cells from tumors were similar to T_EX_ cells in chronic LCMV infection. The 2 migratory cell clusters were dominated by cells from chronic infection, confirming that few tumor-specific CD8 T cells in the dLN or tumor had this distinct gene expression pattern. Notably, cluster 6 (which we classified as a T_SL_ cluster, based on high *Tcf7* expression) had increased expression of many of the migration signature genes, which might suggest that cluster 6 is the closest equivalent to this population in our tumor models. Altogether, these data showed that the tumor-specific CD8 T cell population was a heterogeneous mixture of cells ranging from T_SL_ to T_EX_, and suggested that terminal differentiation may be spatially regulated.

### dLN T_SL_ cells are distinct from memory CD8 T cells

T_SL_ cells share some characteristics with memory T cells (Tmem), so we aimed to directly test whether the T_SL_ populations identified in dLNs were distinct from classical T_mem_. To do this, we compared the transcriptomes between GP33^+^ CD8 T cells from the early and late dLNs to GP33^+^ CD8 T cells sorted from the spleens of mice 28 days after acute LCMV infection (**Figure S3H-J**). The clusters encoding the CD8 T cells from acute infection did not overlap with the clusters encoding the dLN T cells (**Figure S3H**). Moreover, *CCL5* and *Ly6c2* were among the top differentially expressed genes in cells between acute LCMV and early dLN (**Figure S3I**). These genes have been associated with the maintenance (*72*) and homing (*73*) of T_mem_ cells, respectively. By contrast, memory T cells had low expression of the progenitor-and exhaustion-related genes (**Figure 3J**). Together, these data confirm that tumor-specific T_SL_ cells in dLNs were transcriptionally distinct from canonical T_mem_ that are formed after acute infection.

**Figure 3:**
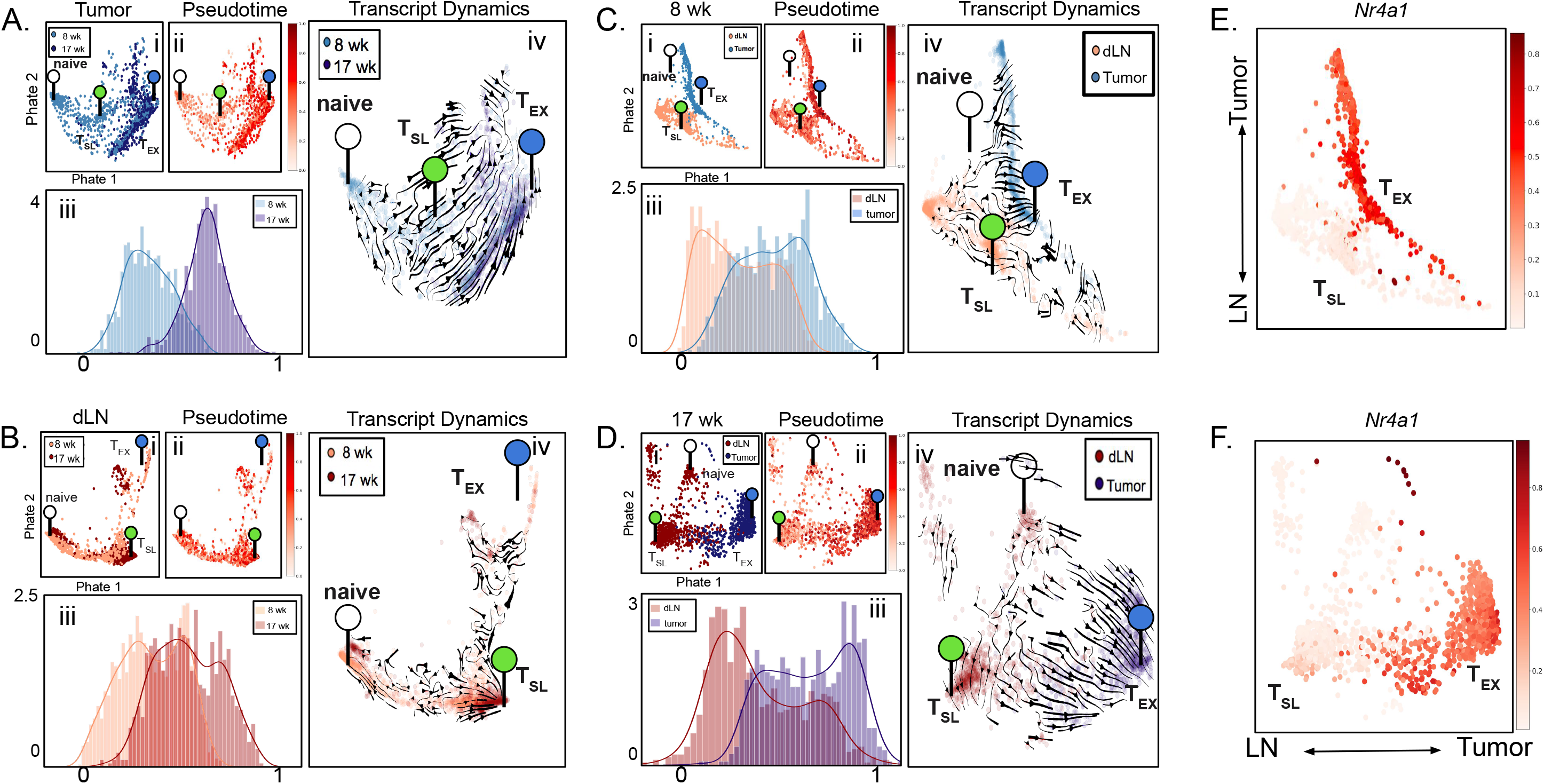
Progressive CD8^+^ T cell differentiation occurs in tumors but not dLNs over the course of disease. (A-D) Single-cell transcriptomics of tumor-specific CD8^+^ T cells from early (8 weeks p.i.) tumors and late (17 weeks p.i.) tumors (A), early dLNs and late dLNs (B), late dLNs and late tumors (C), and early dLNs and early tumors (D) from KP-NINJA mice were co-embedded and expression profiles were visualized by PHATE maps (i). Pseudotime analysis was also visualized by PHATE maps (ii) as well as histograms (iii). Transcript dynamics (iv) between each co-embedded sample pair are illustrated by the direction of arrowheads. The location of transcriptional signatures for the major cell states identified (Naïve (white), T_SL_ (green), and T_EX_ (blue)) are indicated by markers on pseudotime visualizations. (E-F) The gene expression profile for *Nr4a1* visualized by PHATE map on the dLN and tumor co-embeddings at early (E) and late (F) time points.

### Temporal regulation of T-cell differentiation in tumors but not dLNs

To visualize whether CD8 T cell differentiation was regulated at the spatial or temporal level, we co-embedded expression data from GP33-specific CD8^+^ T cells in tumors or dLNs at early or late time points in pairs and visualized their similarities and differences using PHATE. One of the strengths of PHATE is the ability to visualize the transitions in a population without the need to identify discrete clusters of cells. This is ideal for developmental processes that occur along a continuous distribution of differentiation. In each PHATE co-embedding (**Figure 3Ai-Di**), we identified where prototypical naïve (*Ccr7*+ *Sell*+ *Lef*+), T_SL_ (*Tcf7*+ *Xcl1*+ *Slamf6*+), and T_EX_ (*Pdcd1*+ *Havcr2*+ *Cd101*+) cells would lie on the embedding maps (identified with colored markers; see **Figure S4** for gene expression details). This allowed for easier visualization of the transitions between these differentiation states. Moreover, for an unbiased analysis of transitions on our PHATE embeddings we performed pseudotime analyses and leveraged scVelo (*74*), tools optimized to infer trajectories of differentiation using transcriptional splicing kinetics from expression data. Cells were pseudo-colored in each PHATE embedding based on their pseudotime values from 0 to 1 (**Figure 3Aii-Dii**), and associated histograms showed relative cell distribution frequency as a function of pseudotime (**Figure 3Aiii-Diii**). Moreover, a pseudo-temporal ordering of cells from scVelo on the co-embedded tumor samples was extracted and a developmental trajectory was estimated from T_SL_ to T_EX_ (**Figure 3Aiv-Div**). These unbiased analyses supported the idea that in each embedding, the direction of differentiation trended from naïve to T_SL_ to T_EX_. Moreover, this allowed us to determine how tumor-specific CD8 T cell differentiation was regulated by assessing the relative locations of the cells from each sample on the PHATE embeddings.

**Figures 3A and B** show <u>temporal</u> regulation of T cell differentiation in tumors and dLNs, respectively. In the tumor PHATE embedding, we observed distinct naïve and T_EX_ cells, but we had difficulty identifying prototypical T_SL_ cells, consistent with the idea that these cells might not be located in the tumors in our model (**Figure 3A** and **S4A**). The preponderance of T_EX_-like cells were from late tumors, while early tumors contained a mixture of less and more differentiated CD8 T cells. Thus, as tumors developed, we observed a progression of the tumor-specific T cell population towards a more terminally differentiated state. Notably, this corresponded with an overall shift in the TME from T-cell inflamed to non-T cell inflamed between early and late tumors (**Figure 1H**). In dLNs, we identified distinct naïve and T_SL_ cells but had more difficulty discerning a prototypical T_EX_ population (**Figure 3B** and **S4B**). In contrast to tumors, the distribution of cells from early and late dLNs was largely overlapping. In line with this, scVelo analyses confirmed that there was no consistent direction of differentiation between cells from early and late dLNs (**Figure 3Biv**). T cells from dLNs at early and late time points also showed similar distributions across pseudotime (**Figure 3Biii**). The similarity between early and late dLN samples was particularly notable as they were isolated from animals at very different stages of tumor development and their stability stood in sharp contrast to the change observed in T cells in the TME. Together these data demonstrate that over the course of tumor development most tumor-specific CD8 T cells in dLNs remained in a stable, less-differentiated state while the T cells in tumors became progressively more terminally differentiated.

### A continuum of differentiation defines the transition of tumor-specific CD8 T cells from dLNs to tumor

Next we analyzed how spatial location (dLN vs. tumor) impacted tumor-specific CD8 T cell differentiation (**Figure 3C and D**). In both early and late mice, we observed distinct naïve, T_SL_, and T_EX_ cells at both time points, and pseudotime and scVelo analysis supported a continuous differentiation trajectory from the prototypical T_SL_ to the prototypical T_EX_ cells. At both time points, there was a clear segregation of cells from dLNs and tumors, with the less-differentiated T cells being predominantly from dLN and more-differentiated T cells being predominantly from tumors. Yet, in between extremes there was overlap between dLN and tumor, suggesting that the migration of T cells from dLNs to tumors may be an integral component to their differentiation. Pseudotime analysis supported this conclusion, as dLN and tumor T cells were enriched at early and late pseudotime, respectively (0 vs 1, respectively), with more even distribution between (**Figure 3C-D ii and iii**). This provided a means to visualize how the expression of naïve-related, progenitor-related, migration-related, exhaustion-related, effector molecule and transcription factor genes changed as cells differentiated in early and late tumor bearing mice (**Figures S4E-F**). Along the progression of pseudotime (0 -> 1) were decreases in progenitor-related genes and increases in migration-related, exhaustion-related, and effector molecule genes. Moreover, analysis of genes associated with TCR signaling, such as *Nr4a1*, *Nr4a2*, *Nr4a3*, *Egr1*, *Atf3*, *Vps37b*, and *Fosb*, also showed a clear increase across pseudotime (**Figures S4E-F**).

TCR signals are thought to drive terminal differentiation in CD8 T cells responses (*15, 25, 27, 33, 44, 54, 75-78*). To better visualize which T cells were receiving TCR signals, we pseudocolored 8 wk and 17 wk PHATE co-embeddings based on *Nr4a1* (Nur77) expression, a commonly used metric for TCR signaling (**Figure 3G-H**). Strikingly, *Nr4a1* was increased on the more differentiated cells (in tumors), while the T_SL_ cells had *Nr4a1* expression levels similar to naïve T cells. These results were consistent with our earlier analysis of T_SL_ and T_EX_ clusters from dLNs and tumor (**Figure S3B-E**, note TCR signaling genes), and suggest spatial regulation of TCR signals.

### The dLN maintains a reservoir of tumor-specific T_SL_ cells

Despite the above trajectory analysis suggesting that dLN CD8 T cells were differentiating into CD8 T cells in tumors, it remained possible that the T cells in the tumors were maintained independently of those in the dLNs. To assess the lineage relationships between dLN and tumor T cells, we first identified T-cell clones with unique and shared TCR sequences (based on pairs of TCR alpha (TCRA) and TCR beta (TCRB) sequences; **Figure S3F**). We then assessed whether there were shared clones (2 or more cells with the same TCRA and TCRB pair) between early dLNs and tumors and late dLNs and tumors. There was a significant amount of clonal overlap between the GP33-specific CD8 T cells in dLNs and tumors at both early and late time points (**Figure 4A**). Moreover, there was a good correlation between the frequencies of individual clones in the dLN and tumor. Analysis of the top 3 clones in dLN and tumor showed that they were spread across the differentiation trajectory and were present in multiple differentiation states and locations (**Figure 4B**). Moreover, these clonal cells increased in pseudotime value, peaking in the tumors. This demonstrated that tumor-specific CD8 T cells in the dLNs were the developmental precursors of more differentiated intratumoral T cells in mice with early and late tumors.

**Figure 4:**
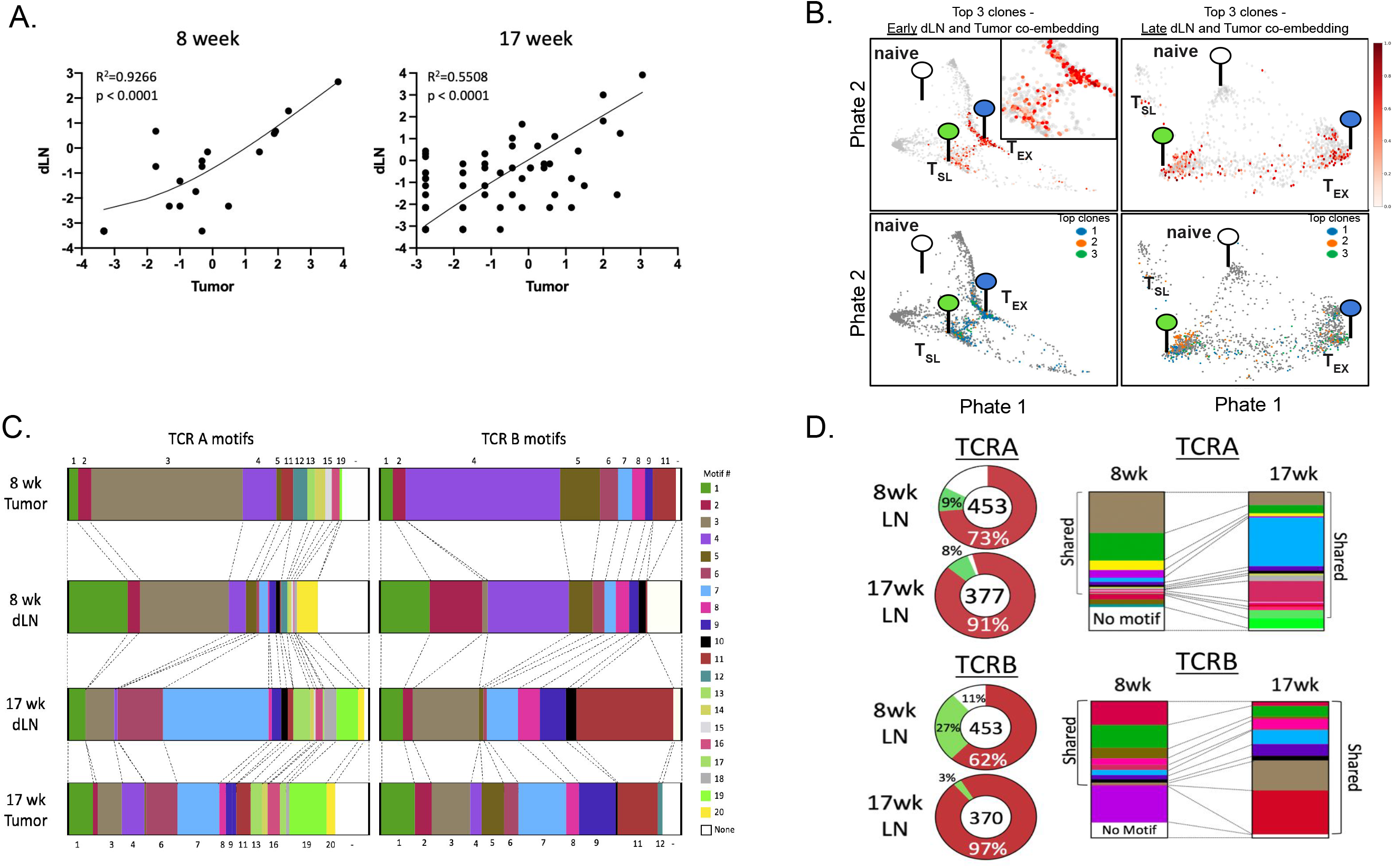
Clonal dominance is maintained throughout disease between dLN and tumors. (A) Paired single cell TCR-sequencing from tumor-specific CD8^+^ T cells (i.v.CD45^-^ CD8^+^GP33-loaded MHC I tetramer^+^) in early (8 weeks p.i.; left) and late (17 weeks p.i.; right) dLNs and tumors was used to identify the correlation between CD8^+^ T cell clones shared between tissues. Abundance of shared each clone is reported as a percent of total shared clones between tissues at either timepoint and displayed on log2 axes (R^2^=0.9226 and 0.5508 at early and late timepoints, respectively). (B) Differentiation status of top 3 shared clones from tumor-specific CD8^+^ T cells in dLN and tumor at early (top left; with area of interest enlarged in top right corner) and late (top right) time points were determined by pseudotime analysis and visualized by PHATE. Distribution of each of the three clones are shown below at early (bottom left) and late (bottom right) time points, with each top clone differently colored. The location of transcriptional signatures for the major cell states identified (Naïve (white), T_SL_ (green), and T_EX_ (blue)) are indicated by markers on pseudotime visualizations. (C) TCRA and TCRB motifs were determined from single-cell TCR-sequencing of tumor specific CD8^+^ T cells and the fraction of clones with each motif are shown. (D) Motifs from early and late dLN displayed in pie charts showing percent “shared” between dLN, “unique” to either early (top) or late (bottom) dLN, or “other” (left) and fraction of individual TCRA motifs (top) and TCRB motifs (bottom) detected in dLNs at either timepoint with portion of shared motifs indicated with brackets (right) and total number of motifs indicated in the center.

The stability of their transcriptional state and the enrichment of early and late dLN T cells amongst the less-differentiated T_SL_ cells raised the question of whether the clones of tumor-specific CD8 T cells in the dLN were maintained over the course of tumor development. Analysis of T cells in the mediastinal LN (tumor-draining) requires sacrifice of animals, so we were unable to directly compare T cell clones between the early and late time points. Thus, we used algorithms that define common TCR motifs for polyclonal populations responding to a common antigen, and we analyzed GP33-specific CD8 T cells from early and late dLNs and tumors (**Figure 4C** and **Table 1**). We identified 20 TCRA motifs and 12 TCRB motifs in total (**Table 1**). As expected, there was good concordance between the presence of TCR motifs in dLNs and tumors of the same mice. Moreover, ∼ 35% of the TCRA motifs and 67% of the TCRB motifs were present in T cells across all four samples, suggesting that T cells with these TCR specificities were maintained over the course of tumor development (referred to here as consensus). Comparing early and late dLN samples, >90% of the 17-wk TCRA and TCRB sequences had motifs that were shared with 8-wk sequences, demonstrating good concordance between T cells in dLNs (**Figure 4D**). By contrast, only 53% of the TCRA sequences and 79% of the TCRB sequences from 17-wk tumors had motifs that were shared with 8-wk sequences (data not shown). These data strongly support the hypothesis that most tumor-specific CD8 T cells in dLNs (and perhaps tumors) are maintained over the course of tumor development.

**Table 1:**
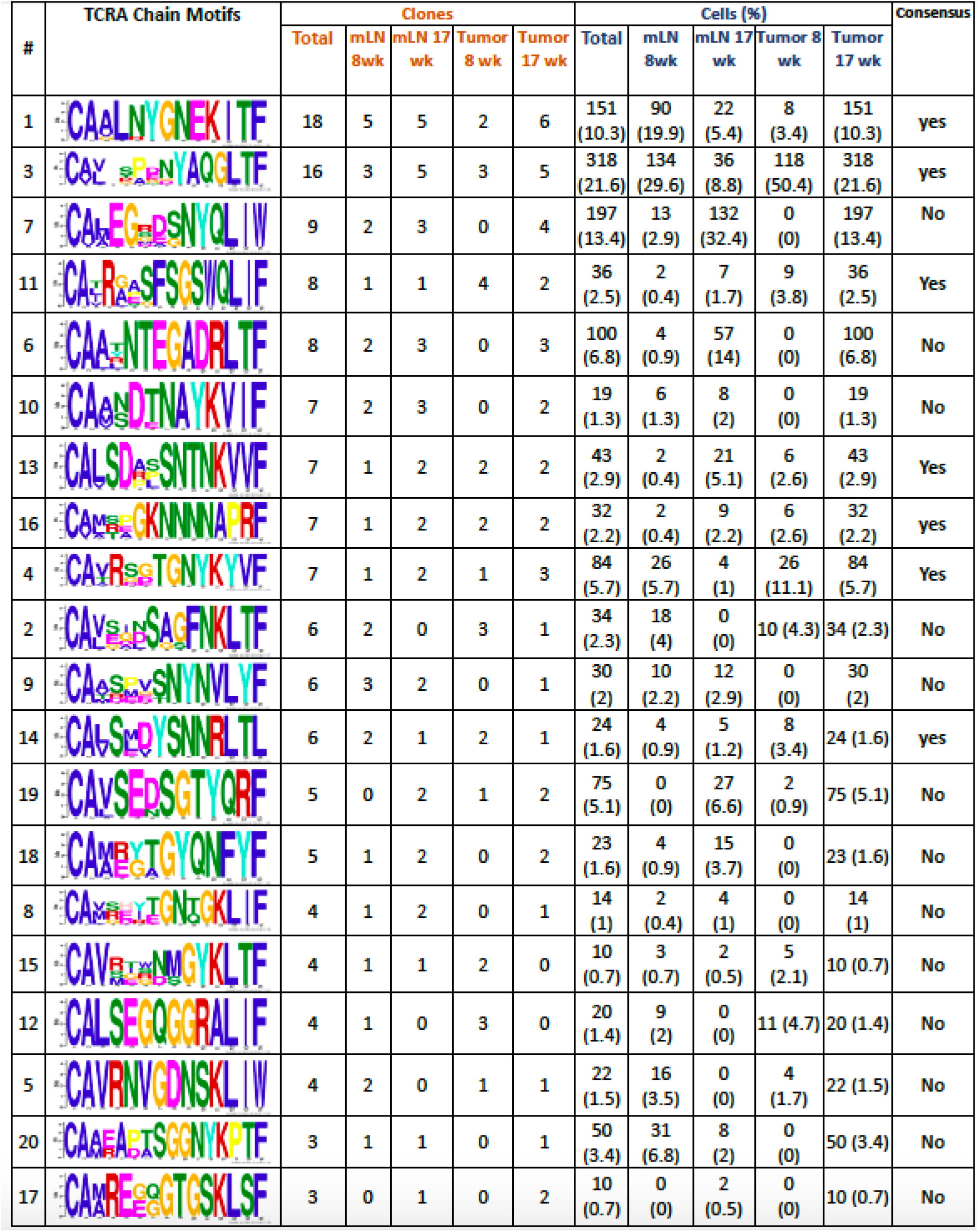
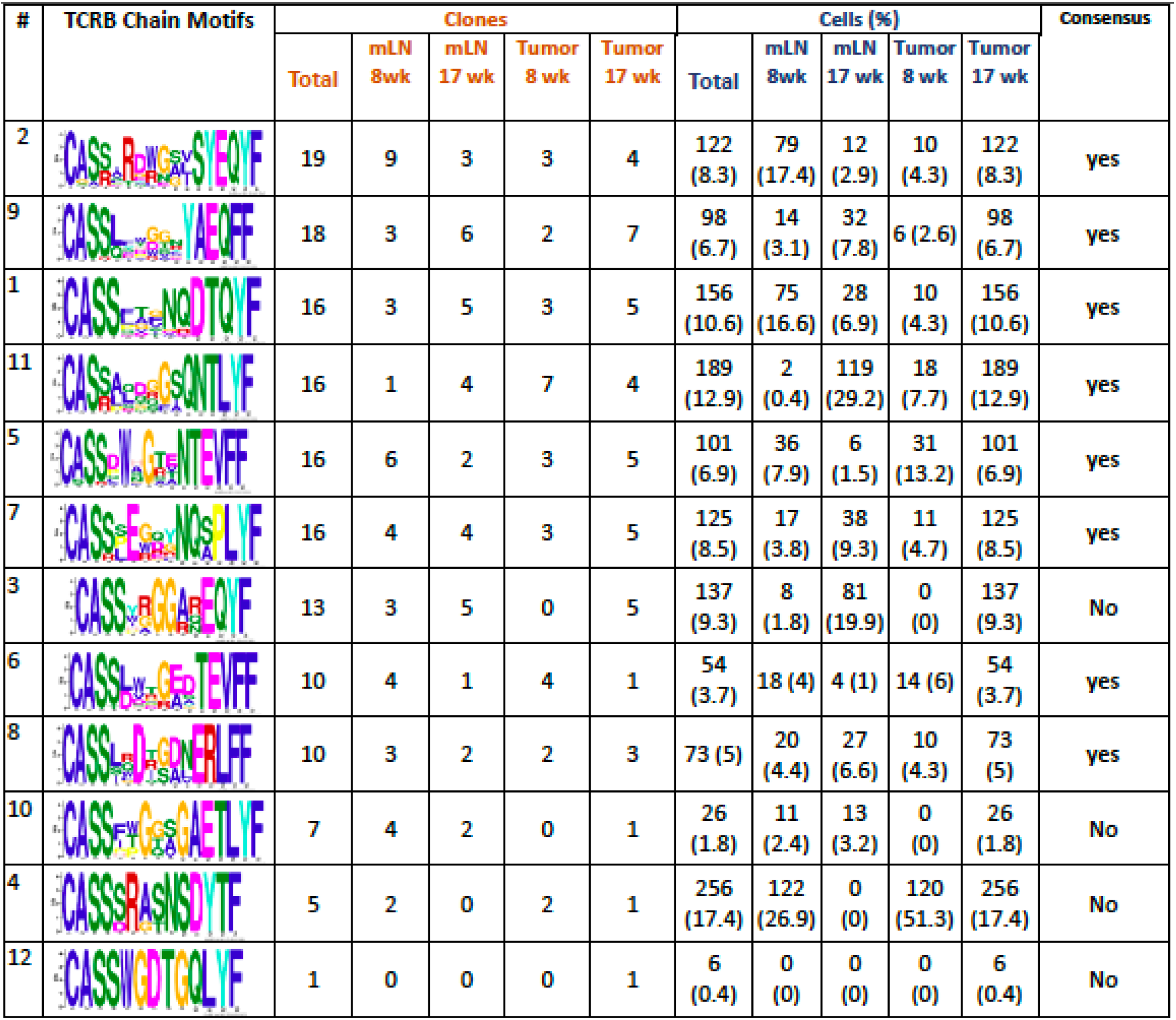
TCRA and TCRB motifs from T cells in tumors and LNs. Consensus motifs in grouped CDR3 amino acid sequences were identified based on single-cell TCR sequencing using two motif based sequencing analysis tools: Multiple Em for Motif Elicitation (MEME) and Gapped Local Alignment of Motifs (GLAM2) (*1*). The motif analysis across all four samples (early and late dLN and tumor) was performed separately for TCR alpha and TCR beta chain, including clones with ≥ 2 cells only. The contribution of clones from each of the four samples to each motif (either for TCR alpha or TCR beta chain), were traced back using the clone IDs. Following this, consensus motif for each chain was defined as having clones shared by all four samples and ranked in an order based on the number of clones giving rise to each motif. The number and percent of clones with shared motifs are reporter for dLN and tumor at early (8 weeks p.i.) and late (17 weeks p.i.)

### Migration from the dLN maintains TCF1 expression by the intratumoral T cell pool

Our data supported two non-mutually exclusive models for the maintenance of T cells in tumors: (1) tumor-specific TCF1^+^ CD8 T cells in tumors could be a self-sustaining population that continued to propagate and differentiate to maintain the T cell response in tumors, or (2) maintenance of TCF1^+^ T cells in tumors could be due to continual migration of small numbers of tumor-specific T_SL_ from the dLN. To test the latter hypothesis, we blocked lymphocyte migration with FTY720 in KP-NINJA mice from 6-9 weeks after tumor induction and assessed the effects on intratumoral tumor-specific CD8 T cells (**Figure 5A**). After 3 wks of FTY720 treatment, the frequency and number of GP33-specific TCF1^+^ CD8 T cells in the tumor tissue was decreased compared to vehicle-treated controls (**Figure 5B-C**). This represented a ∼4-fold drop in the number of these cells (from 1378+/−280 to 363+/−78). By contrast, the number of TCF1^+^ CD8 T cells in the dLN was not impacted by FTY720 treatment, suggesting that the observed decrease was likely due to the impact of blocking migration and not a direct impact of FTY720 on TCF1^+^ CD8 T cells. The observed 4-fold drop in TCF1^+^ CD8 T cells in tumors was more sharp than the 2-fold decrease in the number of total GP33+ CD8 over the same time period in tumors (from 10901+/−2053 to 5750+/−1394), suggesting that the TCF1^+^ T cell population was more impacted by the migration blockade (**Figure 5D-E**). Similar decreases were observed in the GP33-CD8 T cell populations in the tumor (**Figure 5F-G**). Thus, these results are consistent with the hypothesis that the migration of tumor-specific T_SL_ cells from dLNs is necessary to sustain the TCF1 expression of anti-tumor T cells in tumors.

**Figure 5:**
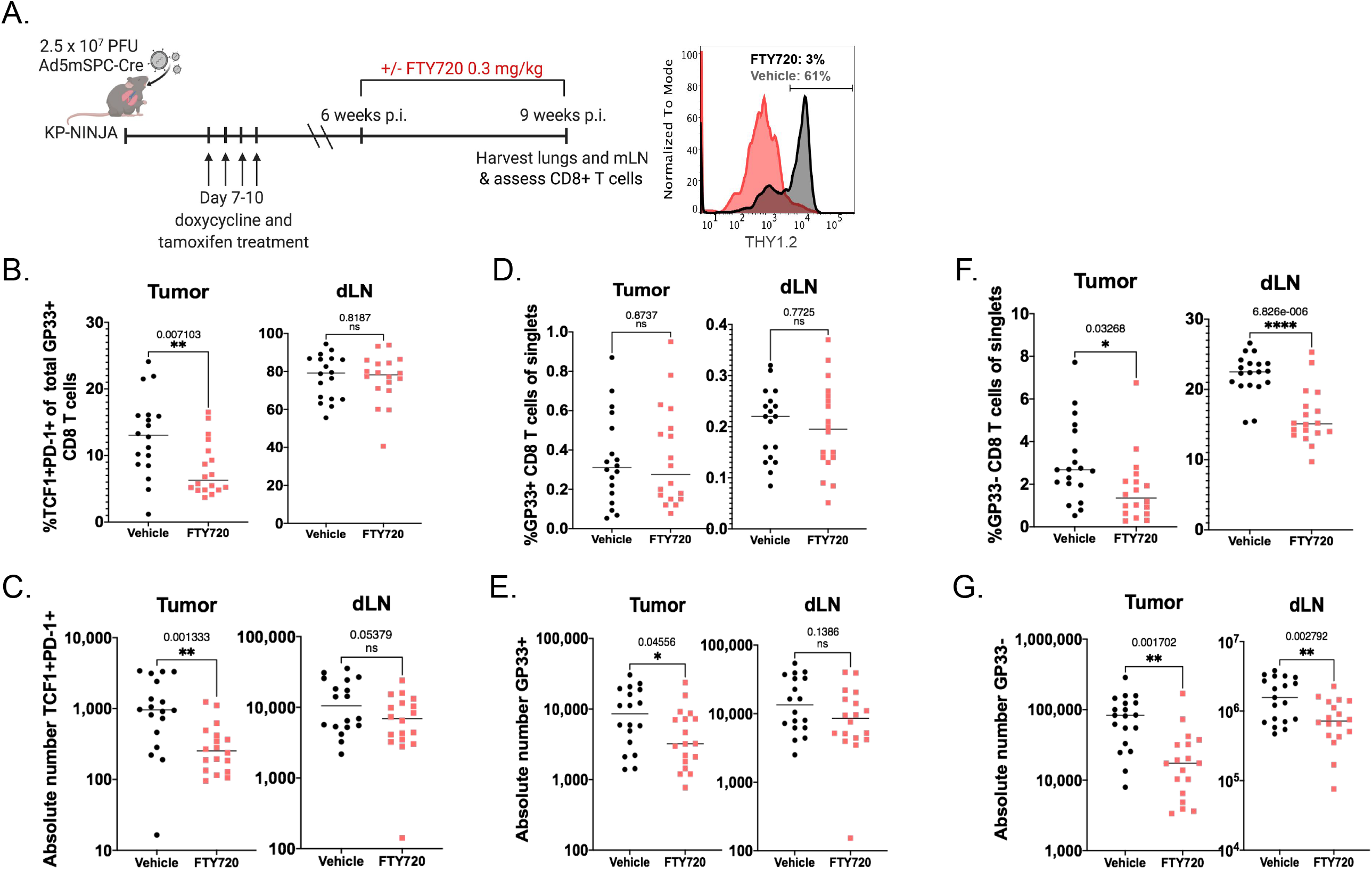
A reservoir of tumor-specific CD8^+^ T_SL_ cells in dLN maintains the anti-tumor immune response. (A) Experimental schematic. Tumors are initiated in KP-NINJA mice via intratracheal infection with 2.5x10^7^ PFU Ad5mSPC-Cre, treated with doxycycline and tamoxifen, then treated with FTY720 or vehicle from 6-9 weeks post-infection and tumor-specific CD8^+^ T cells were assessed at week 9. Representative histogram showing THY1.2^+^ events from whole blood 24hours following 0.3mg/kg FTY720 treatment (gray) and vehicle-treated control (black). (B-C) The percent and absolute number of TCF1^+^ PD-1^+^ T_SL_ in tumors (B, **p=0.007; C, **p=0.001) and dLNs (B, p=0.819; C, p=0.054) in vehicle-treated (black) vs. FTY720-treated (red). (D-E) Perecnt and absolute number of GP-33 specific CD8 T cells in tumors (D, p=874; E, *p=0.046) and dLN (D, p=0.773; E, p=0.139) in vehicle-treated (black) vs. FTY720-treated (red). (F-G) Percent and number of non-GP- 33-specific (GP33-loaded MHC I tetramer-) CD8 T cells in tumors (F, *p=0.033; G, **p=0.002) and dLN (F, ****p=6.8^-6^; G, **p=0.003) in vehicle-treated (black) vs. FTY720- treated (red). Representative of 2 independent experiments containing 4 technical repeats. Statistics based on two tailed, unpaired t-tests; n=18 vehicle and n=18 FTY720 for tumor.

### T_SL_-like populations are present in metastatic and non-metastatic LNs of lung cancer patients

Tumor-draining lymph nodes from human patients are often used for diagnostic purposes and are thus difficult to obtain for research. Therefore, to investigate the potential importance of dLN T cells in human lung adenocarcinoma (LUAD), we took advantage of a recently published single cell RNA sequencing dataset from a study which included normal lung tissue (nLung), tumor-containing lung tissue (tLung), metastatic lymph nodes (mLN), and non-metastatic lung-draining lymph nodes (nLN) collected by surgical resections or ultrasound-guided bronchoscopy biopsies from 44 treatment-naïve LUAD patients (GSE131907; (*79*)). We catalogued 10,046 CD8 T cells from all four tissues, based on their previous annotations, and these were grouped into 15 clusters and visualized using the dimension reduction method Uniform Manifold Approximation Projection (UMAP) (**Figure 6A**).

**Figure 6:**
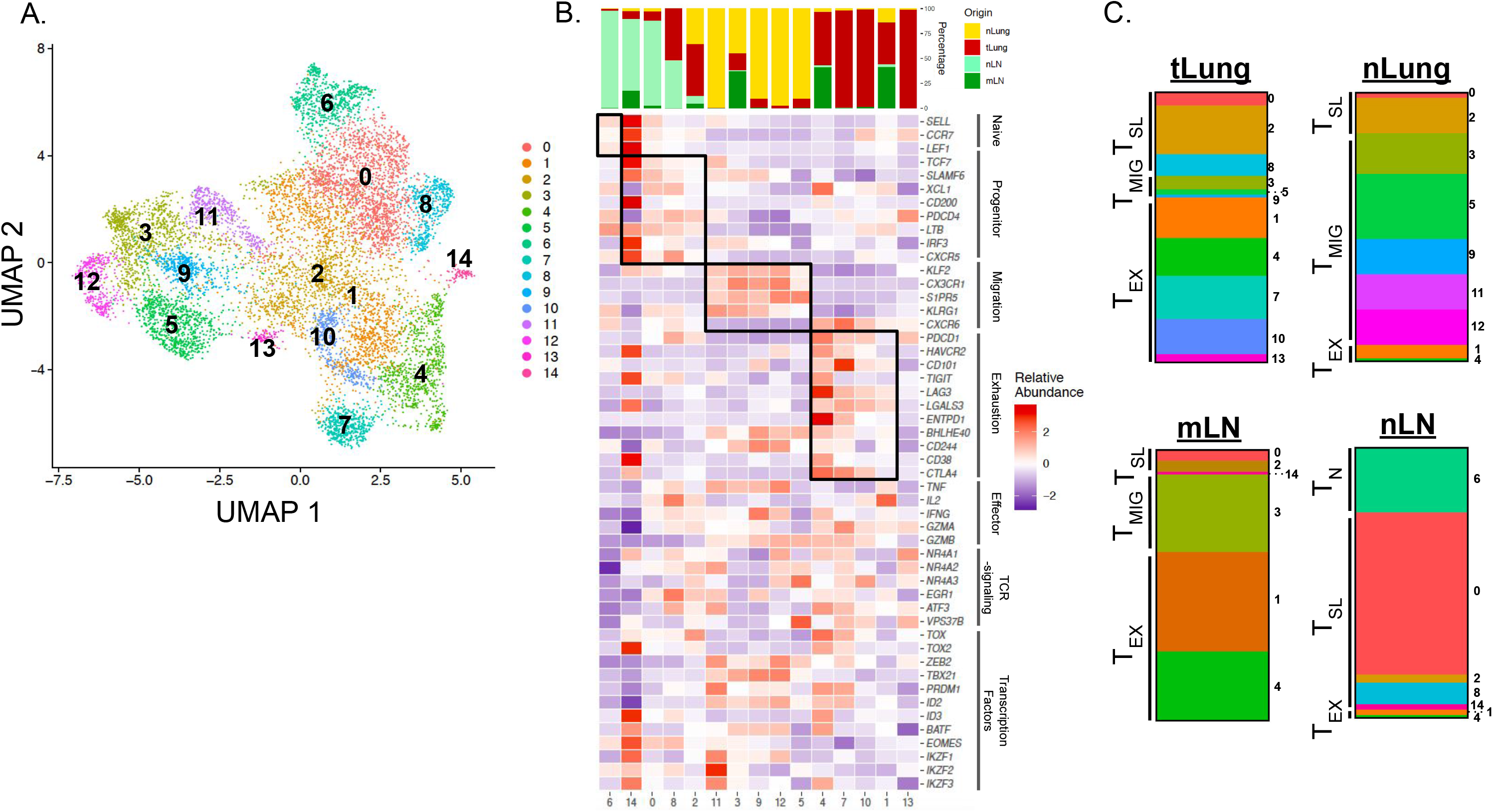
T_SL_-like populations are prevalent in non-metastatic LN of lung cancer patients. (A) UMAP displaying CD8 T cell clusters 0-14 in treatment naïve LUAD patients from primary sites (tLung) and metastatic LN (mLN), normal lung tissue (nLung) and non- metastatic, normal LNs (nLN) (GSE131907; (*79*)). (B) Tissue of origin distributions for each cluster (top) and heatmap displaying relative abundance for naïve related (*Sell, Ccr7, Lef1*), progenitor-related (*Tcf7, SLAMF6, Xcl1, Cd200, Pdcd4, Ltb, Irf3, Cxcr5*), migration-related (*Klf2, Cx3cr1, S1pr5, Klrg1, Cxcr6*), and exhaustion-related (*Pdcd1, Havcr2, Cd101, Tigit, Lag3, Lgals3, Entpd1, Bhlhe40, Cd244, Cd38, Ctla4*) signature genes (bottom). Key effector cell-associated genes (*Tnf, Il2, Ifng, Gzma, Gzmb*), genes associated with intracellular TCR signaling (*Nr4a1, Nr4a2, Nr4a3, Egr1, Atf3, Vps37b*), and key transcription factors (*Tox, Tox2, Zeb2, Tbx21, Prdm1, Id2, Id3, Batf, Eomes, Ikzf1, Ikzf2, Ikzf3*) are also included. Relative abundance was calculated as *Z*-scaled average of log-transformed and cell-normalized counts. (C) Bar graphs depicting the make-up of total CD8 T cells from each tissue showing the distribution of T_N_-like, T_SL_-Like, T_MIG_-like, or T_EX_-like clusters determined by gene signatures from B.

With the same signature genes used in **Figure 1**, we grouped CD8 T cell clusters from LUAD patients into naïve T cell (T_N_), T_SL_-like, migratory T cell (T_MIG_)-like, and T_EX_-like categories (**Figure 6B**). We identified 1 naïve cluster (1), 4 clusters with a T_SL_-like signature (14, 0, 8, 2; ordered from highest to lowest *TCF7* expression), 5 clusters with a T_MIG_-like signature (11, 3, 9, 12, 5; ordered from highest to lowest *KLRG1* expression), and 4 clusters with a T_EX_-like signature (4, 7, 10, 1; ordered from highest to lowest *PDCD1* expression). Gene expression signatures of T_SL_-like and T_EX_-like clusters were strikingly similar to these populations from tumor-bearing KP-NINJA mice in our model (refer to **Figure 2F**). Furthermore, we were struck by the tissue distributions of the clusters within these categories, as 73% of the nLN CD8 T cells were T_SL_-like (**Figure 6C**), in agreement with our findings in mice. T_SL_-like cells made up 31% of CD8 T cells from the tLung, and only 9% and 15% of the CD8 T cells from the mLN and nLung, respectively. In contrast, T_EX_-like cell clusters were dominant in the mLN and tLung (61% and 61%, respectively). Interestingly, nLung tissue contained cells which belonged predominantly to T_MIG_-like clusters (78%). Together, these data from LUAD patients support our hypothesis that T_SL_ cells reside in the dLNs and that differentiation occurs after the dLN T cells migrate to the tumor.

## Discussion

PD-1^+^ TCF1^+^ CD8 T cells are present in human and murine tumors and are necessary to sustain both the anti-tumor T cell response and responses after immunotherapy, but the mechanisms for their maintenance remained unclear. Using an autochthonous model of lung adenocarcinoma, we found that the population of intratumoral PD-1^+^ TCF1^+^ CD8 T cells was maintained by migration from the tumor-draining lymph node (dLN). Most tumor-specific CD8 T cells in dLNs expressed PD-1, TCF1, and SLAMF6 and had transcriptional patterns that were more similar to canonical T_SL_ cells seen during chronic LCMV infection. By contrast, while some intratumoral CD8 T cells were PD-1^+^ and TCF1^+^, many did not express SLAMF6 or other genes associated with T_SL_ cells. Moreover, as the tumor microenvironment shifted from hot to cold, the intratumoral T cell population became more differentiated, while the population in the dLN was unchanged at the transcriptional and phenotypic levels. dLN T cell TCR motifs were also maintained over several months. Thus, while most studies of CD8 T cell function and differentiation have focused on tumor tissues (*80–88*), our data demonstrate that the process of differentiation for tumor-specific CD8 T cell begins in the dLN, with the dLN serving as a reservoir for maintaining T cells in a stem-like state throughout the course of tumor development.

The question of whether patients with cold tumors can respond to immunotherapeutic intervention has remained uncertain. Cold tumors have poor infiltration of T cells and/or T cell exclusion, which is thought to reflect a diminished or absent anti-tumor immune response, consistent with their poor response to checkpoint therapies (*4*). Yet, cold tumors have similar mutational burdens and antigen presentation capacity as hot tumors, suggesting they both have the capacity to initiate and drive anti-tumor T cell responses (*53*). These findings are in line with the cold tumors in our model, which maintain neoantigen expression and *in vivo* presentation of neoantigens (*49, 60, 89, 90*). Given these observations, we hypothesized that the cold tumor microenvironment was a result of global exhaustion of tumor specific CD8 T cells throughout the host. Surprisingly, many tumor-specific CD8 T cells in late tumors were TCF1^+^, although pseudotime analyses demonstrated that these T cells were more differentiated than T cells from early tumors. These data are consistent with the possibility that the increased differentiation state of intratumoral T cells could account for the cold phenotype of late tumors. Additional possibilities include the inability of migrating T cells to physically enter tumors or defects in DC migration or function (*53, 91–93*). Yet, we also found that a cold TME is not indicative of global exhaustion of tumor-specific CD8 T cells as cold tumors were associated dLNs containing T_SL_ cells that were transcriptionally similar to T_SL_ cells in dLNs associated with hot tumors. Thus, the distal location of dLN T_SL_ cells likely protects them from changes that occur within the TME over the course of tumor development.

While many signals could promote the differentiation of intratumoral CD8 T cells, TCR signals are prime candidates. TCR signals drive terminal T cell differentiation in chronic infection, and CD8 T cells that recognize more abundant antigens are subject to more severe exhaustion (*15, 25, 27, 33, 44, 54, 75-78*). Likewise, antigenic peptides that deliver weaker TCR signals are less potent drivers of T cell exhaustion (*44*). *Tcf7* is required to sustain CD8 T cells during chronic infection (*31, 40, 44*), but TCR and inflammatory signals promote TCF1 downregulation (*94*). We found that intratumoral T cells had high levels of transcripts associated with downstream TCR signaling, while dLN T cells had low expression of these transcripts. These data are consistent with the idea that the dLN protects T_SL_ cells from persistent antigen exposure. We hypothesize that intratumoral CD8 T cells are unable to escape persistent antigen and that without migration from the dLN, the pool of tumor-specific CD8 T cells would become exhausted. Moreover, because T cell clones that recognize tumor antigens with higher avidity (so called “best fit” clones) are more prone to exhaustion, the dLN likely plays an important role in preventing the loss of best fit clones over the course of tumor development. This is consistent with our data showing the maintenance of TCR motifs in the tumor and dLN between 8 and 17 wks. The splenic white pulp may play a similar role during chronic LCMV infection (*31, 38*), although both the white and red pulps are sites of LCMV Clone 13 infection (*95*). Our data also raise the question of whether intratumoral niches (like TLSs) could exist in tumors to protect T_SL_ cells from differentiation. We previously showed that TLSs associated with late tumors in our models were sites for antigen presentation (*60*), but it remains to be seen whether tumor-proximal niches such as TLSs could protect resident T cells from persistent antigen exposure (*60, 89, 90*).

Our data highlight the critical role of migration in the maintenance of T_SL_ cells in tumors but are less consistent with the idea that T cells differentiate in the lymphoid tissue prior to migration. The latter has been seen in chronic LCMV infection (*31, 38*) and may be due to the ongoing infection in the tissue. By contrast, we observed T_SL_ cells differentiate upon migration to tumors, and that migration was required for the presence of less differentiated T cells in tumors. It is not clear what drives the migration of T_SL_ cells from dLNs to tumors. DC migration from tumors to dLNs is important for priming T cells, but the role of DCs in maintenance of T_SL_ cells in dLNs is uncertain (*96, 97*). One simple model is that periodic signals from migrating DCs are also necessary for maintaining the migratory T cell population. Critically, while we did not see the accumulation of less-differentiated T cells in dLNs upon FTY720 treatment, it is possible that FTY720 also blocks the migration of antigen-presenting DCs to LNs, which could impact the differentiation of T_SL_ cells in the dLN. Further studies will be needed to test what signals are necessary for maintenance and migration of T cells in dLNs.

The role of the dLN in immunotherapy remains uncertain. Expression of PD-L1 on DCs is important for responses to anti-PD-L1 in some tumor models, and migratory DCs in tumor dLNs express both PD-L1 and the costimulatory receptor B7-2 (*98*). Moreover, PD-1 blockade can act in dLNs in transplant tumor models (*99, 100*). Whether PD-1 blockade acts outside the TME in humans is not known, but therapeutic efficacy after anti-PD-1 treatment in patients is associated with changes in immune cell populations in the peripheral blood (*101, 102*) and with the appearance of new T cell clones in the tumor after therapy (*103–105*). Our analyses of CD8 T cells from humans showed the presence of T_SL_-like cells in LNs and tumors. However, as PD-1 blockade functions poorly in patients with cold tumors, this suggests that these patients either lack LN T_SL_ cells or that PD-1 blockade is insufficient to drive therapeutic responses in LNs of these patients. Here, it is notable that tumor-specific T_SL_ cells express a number additional therapeutic targets, including several co-stimulatory receptors like OX-40 and 4-1BB (data not shown). Thus, identifying novel therapeutic strategies directed towards tumor-specific T cells in the dLN may be a means towards improving outcomes for cancer patients with cold tumors.

## Materials and Methods

### Mice

C57BL/6J mice (Jackson Laboratories) were used for all transplant experiments. KP x CCSP-rtTA mice, referred to here as KP mice were obtained from Tyler Jacks lab (*50*) and crossed to NINJA mice (*59*) to obtain KP-NINJA (Kras^lslG12D/+^, p53^fl/fl^, R26-NINJA/NINJA, CCSP-rtTA^+^) mice. KP (Kras^lslG12D/+^, p53^fl/fl^, CCSP-rtTA^+^) mice were used as controls in some cases. 6+ week-old male and female adult mice were used for all experiments. All studies were carried out in accordance with procedures approved by the Institutional Animal Care and Use Committees of Yale University. All mice were bred in specific pathogen-free conditions.

### Lung Tumor Initiation

Autochthonous tumor generation: KP-NINJA mice were infected intratracheally with 2.5 x 10^7^ PFU Ad5mSPC-Cre (Dr. Anton Berns, Netherlands Cancer Institute), after precipitation with 10mM CaCl_2_ for 20-60 minutes, or 5 x 10^4^ PFU Lenti-cre. To induce expression of NINJA neoantigen in infected cells, mice were given doxycycline hyclate chow (625mg/kg; Envigo cat. TD.09628) days 7-11 post infection (p.i.) and concomitantly treated with 4.4mg tamoxifen (MP Biomedicals cat. MP215673894) in corn oil (ThermoFisher Scientific cat. S25271) by gavage on days 8-10 p.i. To induce neoantigen expression via lentivirus, KP mice were infected with 2.5 x 10^4^ PFU mClover-GP33-80-Cre lentivirus and assessed at 8 weeks pi. Orthotopic KPN1 tumor transplants: Established KPN1 cells were maintained in complete DMEM (10% HI-FBS, 55μM beta-mercaptoethanol, 1x Pen/Strep and 1x L-Glut). Prior to injection, cells were washed 3x with 1xPBS and 200,000 cells were injected intravenously via tail vein injection. Subcutaneous KPN1 transplants: Established KPN1 cells, sorted for GFP+ (NINJA-expressing) cells, were maintained in complete DMEM (10% HI-FBS, 55μM beta-mercaptoethanol, 1x Pen/Strep and 1x L-Glut). Prior to injection, cells were washed 3x with 1xPBS and 500,000 cells were injected s.c. and measured using standard caliper measurements. Tumor volume = (LxW^2^)/2.

### Lung and LN processing for flow cytometry

Prior to sacrifice, mice were injected retro-orbitally with 200uL anti-CD45-PECF594 in 1X PBS (1:200; BD Biosciences Cat# 562420). After 2-3 minutes, lungs were harvested at various time points ranging from 8-25 weeks p.i. into Collagenase IV (Worthington Biochemical, cat. LS004189) Buffer (1x HEPES buffer, 0.5mg/mL Collagenase IV, 20μg/mL DNase in 1x HBSS with MgCl_2_ and CaCl_2_) and run on the default Lung_01 protocol on a gentleMACS Dissociator instrument (Miltenyi Biotec). Samples were then incubated at 37°C for 30 min and further dissociated with default Lung_02 protocol. Digestion was quenched by adding 500 μL FBS. Tumor samples were then strained through 70 μm cell strainers, washed with 1% HI-FBS RPMI-1640 (ThermoFisher Scientific cat. 11875085) and red blood cells were lysed using 1x RBC Lysis Buffer (eBioscience, cat. 00-4333-57). Cells were counted using a hemocytometer for absolute number calculations. Mediastinal lymph nodes were concomitantly harvested from all tumor-bearing mice, and processed as described in (*106*). Single cell suspensions were stained using one of two antibody panels (see tables) in addition to tetramer for H2Db/GP_33-43_-specific CD8 T cells (NIH Tetramer Core Facility). For intracellular staining, FoxP3/Transcription Factor Staining Buffer set (eBioscience cat# 00-5523-00) was used as per manufacturer’s protocol. Cells were washed and resuspended in FACs Buffer (0.5% FBS, 20% sodium azide in water, PBS 1X without Mg^2+^/Ca^2+^) until analysis on a BD LSRII flow cytometer (BD Biosciences).

### *Ex Vivo* IFNγ expression

Single cell suspensions were obtained as described above. The number of cells from draining lymph nodes and tumors was determined using hemocytometer. Samples were then plated in 96-well flat bottom plates at a ratio of 25:75 with CD45.1 splenocytes and stimulated in 10% HI-FBS RPMI-1640 (ThermoFisher Scientific cat. 11875085) containing Brefeldin A (eBioscience cat. 00-4506-51), and LCMV GP_33-41_ peptide (AnaSpec cat. AS-61296), or left unstimulated in 10% HI-FBS RPMI-1640 (ThermoFisher Scientific cat. 11875085) containing Brefeldin A (eBioscience cat. 00-4506-51). Plates were incubated for 4-6 hours at 37°C, and samples were then transferred to 96-well round bottom plates. Samples were then stained for extracellular markers (see tables), fixed with BD Cytofix/Cytoperm Fixation/Permeabilization Solution kit (BD Biosciences cat. 554714), and stained with anti-IFNγ for intracellular cytokine assessment (see tables) in BD Perm/wash Buffer (BD Biosciences cat. 554714) as per manufacturer’s protocol.

### Histology and IHC staining

Tumor-bearing lungs of KP-NINJA mice were fixed in 1x Formalin solutions in PBS (Millipore-Sigma) for 24 hours at 4°C, switched into 70% ETOH, and submitted to Yale histology core for paraffin embedding, sectioning, and hematoxylin and eosin (H&E) staining. Unstained slides of KP-NINJA autochthonous lung tumors were stained with anti-CD3 (ab5690) using the ImmPACT DAB Peroxidase kit (Vector Labs) for immunohistochemistry. H&E and anti-CD3 IHC stained sections were imaged on a Nikon TE2000 microscope (Micro Video Instruments, Inc. Avon, MA) using a 20x objective.

### FTY720 Treatment

KP-NINJA mice were infected intratracheally with 2.5 x 10^7^ PFU Ad5mSPC-Cre (Dr. Anton Berns, Netherlands Cancer Institute) and treated with tamoxifen and doxycycline as previously described. From 6 to 9 weeks following intratracheal infection, mice were treated with 0.3 mg/kg FTY720 or vehicle (saline) i.p. every other day.

### Cell line generation

KP-NINJA mice were infected intratracheally with 5 x 10^4^ PFU of lentiviral vector LV-rtta3-Cre and KPR mice with LV-LucOS-Cre. KPRN mice were treated with doxycycline and tamoxifen as described above to induce NINJA expression in transformed cells. Lungs of all mice were harvested 20 weeks p.i., minced with scissors, and rotated at 37 C for 40 minutes in Collagenase IV Buffer + 2 mg/mL Dispase II (Sigma Aldrich cat. 04942078001). Homogenate was filtered through a cell strainer (Corning cat. 352340) and centrifuged at 200xg for 4 minutes at room temperature. Pellet was resuspended and cultured at 37°C and 5% CO_2_ in complete DMEM (DMEM + 10% FBS + 1% P/S), + 1x Gentamicin for the first 2 passages. After 6+ passages fibroblasts were visually undetectable and cell lines were verified to be 100% Kras-transformed by treating with puromycin (unrecombined kras in this mouse confers puromycin resistance, data not shown).

### LCMV-Clone 13 and -Armstrong infections

For Chronic and acute LCMV infections, 7-10 weeks old C57BL/6 mice were infected intraperitoneally with 2x10^6^ PFU/mouse of LCMV-Clone 13 or LCMV-Armstrong, respectfully. Mice were euthanized 28 days after infection to collect and process spleens as previously described (*106*).

### Sorting and single cell RNA- and TCR-sequencing of tumor-specific CD8 T cells

KP-NINJA mice were infected with Ad5mSPC-Cre, treated with doxycycline and tamoxifen, and lungs and dLN were harvested 8 and 17 weeks p.i. after i.v. injection of anti-CD45-PECF594 antibody (clone 30-F11, BD Biosciences), as described above. Tissues were dissociated as previously described and single cells were sorted on intravascular CD45^-^CD8 GP33-loaded MHC I tetramer^+^ events. Data represents cells from n=3 pooled at each time point. Pooled GP33-specific endogenous cells from tumors and matched dLNs were submitted for 10X single cell RNA-and TCR-sequencing to Yale Center for Genome Analysis (YCGA).

### Motif Analysis

Consensus motifs in grouped CDR3 amino acid sequences were identified using two motif based sequencing analysis tools: Multiple Em for Motif Elicitation (MEME) and Gapped Local Alignment of Motifs (GLAM2) (*1*). The motif analysis across all four samples (early and late dLN and tumor) was performed separately for TCR alpha and TCR beta chain, including clones with ≥ 2 cells only. At first, fasta files were created separately, including TCR alpha or TCR beta CDR3 amino acid sequences for the clones with ≥ 2 cells, using Biostrings package. These fasta files were used as input files for motif analysis, separately for each chain. Filtration of CDR3 amino acid sequences were performed based on low alignment scores by GLAM2. From there, CDR3 amino acid sequences for each chain were further sub-grouped into separate fasta files based on similarity in sequences and alignment scores. Each sub-group of sequences for each chain were run for motif analysis using GLAM2 function and a position weight matrix as an output to define the motif for each sub group of sequences, either for TCR alpha or beta chain. The contribution of clones from each of the four samples to each motif (either for TCR alpha or TCR beta chain), were traced back using the clone IDs. Following this, consensus motif for each chain was defined as having clones shared by all four samples and ranked in an order based on the number of clones giving rise to each motif (Table 1).

### Bioinformatics analysis of GP33-specific CD8 T cells

Single cell RNA- and TCR-sequencing data from LCMV-Clone 13, LCMV-Armstrong, early and late dLNs and tumors was processed with CellRanger 3.1 using the mm10 mouse genome indices from 10x genomics. Number of cells analyzed and genes detected for each sample: Chronic LCMV(1,185 and11,595), Acute LCMV(10,768 and 12,960), early dLN (1,742 and 12,116), late dLN (876 and 11,595), early tumor (806 and 11,749), and late tumor (731 and 11,150). The libraries were further pre-processed in Python using the scprep package (github.com/krishnaswamylab/scprep). Cells with library size below 1000 UMI/cell and rare genes (genes detected in fewer than 5 cells) were removed. The data was then normalized by library size to 1,000 counts per cell and square-root transformed.

For visualization, PHATE (*64*) was used to embed the cells into two dimensions based on transcriptional profiles, allowing for visual comparison of global and local similarities between cells. Groups of similar cells were identified by running spectral clustering on our input data. For visualizing gene expression, we imputed missing and dropped-out values with MAGIC and visualized on PHATE (*107*). Finally, a small percentage of the cells were found to have low expression of *CD8a* after de-noising were excluded from further analysis.

To analyze the cellular trajectories and infer pseudotime, we used the scVelo stochastic model (*74, 108*) stochastic model. Pseudotime was computed on the basis of the inferred velocity graph with scVelo.

### Human CD8 T cell single cell RNA-sequencing analysis

Single cell data was obtained from Gene Expression Omnibus (GEO) by accession code GSE131907 (*79*). Data processing, analysis and visualization were conducted using R program with package Seurat (v 3.1.0). Only CD8+ T cells (original labels “CD8 low T”, “Cytotoxic CD8+ T”, “Naive CD8+ T” and “Exhausted CD8+ T”) with tissues from tumor or normal lungs, and metastatic or normal lymph nodes (original labels “tLung”, “nLung”, “mLN” and “nLN”) were used for the analysis. Raw gene counts were log-normalized by Seurat function NormalizeData with parameter normalization.method set to “LogNormalize”. Cell clusters were identified from the normalized data using functions FindNeighbors and FindClusters on top 20 PCs and resolution 0.75. UMAP was used to visualize cell clusters based on top 20 PCs. For the marker gene expression heatmap, relative abundances for each gene were calculated as Z-scaled average of log2(RC+1). Here RC are relative counts calculated by Seurat function NormalizeData with parameter normalization.method set to “RC”.

### Statistical analyses

All statistical analyses were performed using Prism V8.3.0 software.

### Figure Design

Figures 1A, 1B, 2A, and 5A were created with BioRender.

| Antibody Panel 1 | Source | Identifier | Dilution |
| --- | --- | --- | --- |
| CD45-PECF594<br>(clone:30-F11) | BD Biosciences | Cat# 562420 | 1:200 |
| CD90.2-PERCP<br>(clone:30-H12) | BioLegend | Cat# 105322 | 1:500 |
| CD8a-FITC<br>(Clone:CT-CD8a) | Thermo Fisher Scientific | Cat# MA5-17598 | 1:500 |
| CD279-BV421<br>(clone:29F.1A12) | BioLegend | Cat# 135217 | 1:500 |
| TCF1/TCF7-PE<br>(Clone:C63D9) | Cell Signaling<br>Technologies | Cat# 14456S | 1:500 |

| Antibody Panel 2 | Source | Identifier | Dilution |
| --- | --- | --- | --- |
| CD45-PECF594<br>(clone:30-F11) | BD Biosciences | Cat# 562420 | 1:200 |
| CD8-PECy5<br>(clone:CT-CD8a) | Thermo Fisher Scientific | Cat# MA5-17601 | 1:500 |
| Thy1.1-APCFire750<br>(clone:OX-7) | BioLegend | Cat# 202544 | 1:500 |
| Thy1.2-BV605<br>(clone:30-H12) | BioLegend | Cat# 105343 | 1:500 |
| Tim3-PECy7<br>(clone:RMT3-23) | BioLegend | Cat# 119716 | 1:500 |
| PD-1-BV421<br>(clone:29F.1A12) | BioLegend | Cat# 135217 | 1:500 |
| SLAMF6-PE<br>(clone:330-AJ) | BioLegend | Cat# 134606 | 1:500 |

| Peptide re-stimulation panel | Source | Identifier | Dilution |
| --- | --- | --- | --- |
| CD8α-BV421<br>(clone:53-6.7) | BioLegend | Cat# 100737 | 1:500 |
| CD45.1-APC-Cy7<br>(clone:A20) | BioLegend | Cat# 110716 | 1:500 |
| Thy1.2-BV605<br>(clone:30-H12) | BioLegend | Cat# 105343 | 1:500 |
| IFNγ-FITC<br>(clone:XMG1.2) | BD Biosciences | Cat# 55441 | 1:200 |

## Acknowledgements

We thank Joshi lab members for reviewing the manuscript. We also thank the Yale Cancer Center (P30 CA016359 40), Yale Flow Cytometry Core, Yale Center for Genomics Analysis, and Yale School of Medicine Histology Facility. For Ad5mSPC-Cre we thank Dr. Anton Berns of the Netherlands Cancer Institute. We also thank Dr. John Wherry (University of Pennsylvania) for the generous gift of LCMV clone 13. This work was supported by grants from the NCI K22CA200912 (N.S.J.), Young Investigator Award-Melanoma Research Alliance (N.S.J.), Career Enhancement Award from Yale SPORE in lung cancer 1P50CA196530 (N.S.J.), a grant from the Lung Cancer Research Foundation (LCRF) (N.S.J.), NCI 1RO1CA237037-01A1 (N.S.J.), an American Lung Association Discovery Award (N.S.J.), the Yale Cancer Center Leslie Warner Postdoctoral Fellowship (K.A.C.), the Interdisciplinary Immunology Training Program: grant number: NIH AI07019 (K.A.C.), AI125741 (W.C.), AI148403 (W.C.), American Cancer Society Research Scholar Grant (W.C.). M.Y.K. is a member of the Medical Scientist Training Program at the Medical College of Wisconsin, which is partially supported by a training grant from NIGMS (T32-GM080202). This work was also funded in part by the NHLBI-funded postdoctoral fellowship: T32 HL007974 (G.F.). GF is a PhD Student in the Investigative Medicine Program at Yale which is supported by CTSA Grant Number UL1 TR001863 from the National Center for Advancing Translational Science (NCATS), a component of the National Institutes of Health (NIH).

---

**Figure S1:**
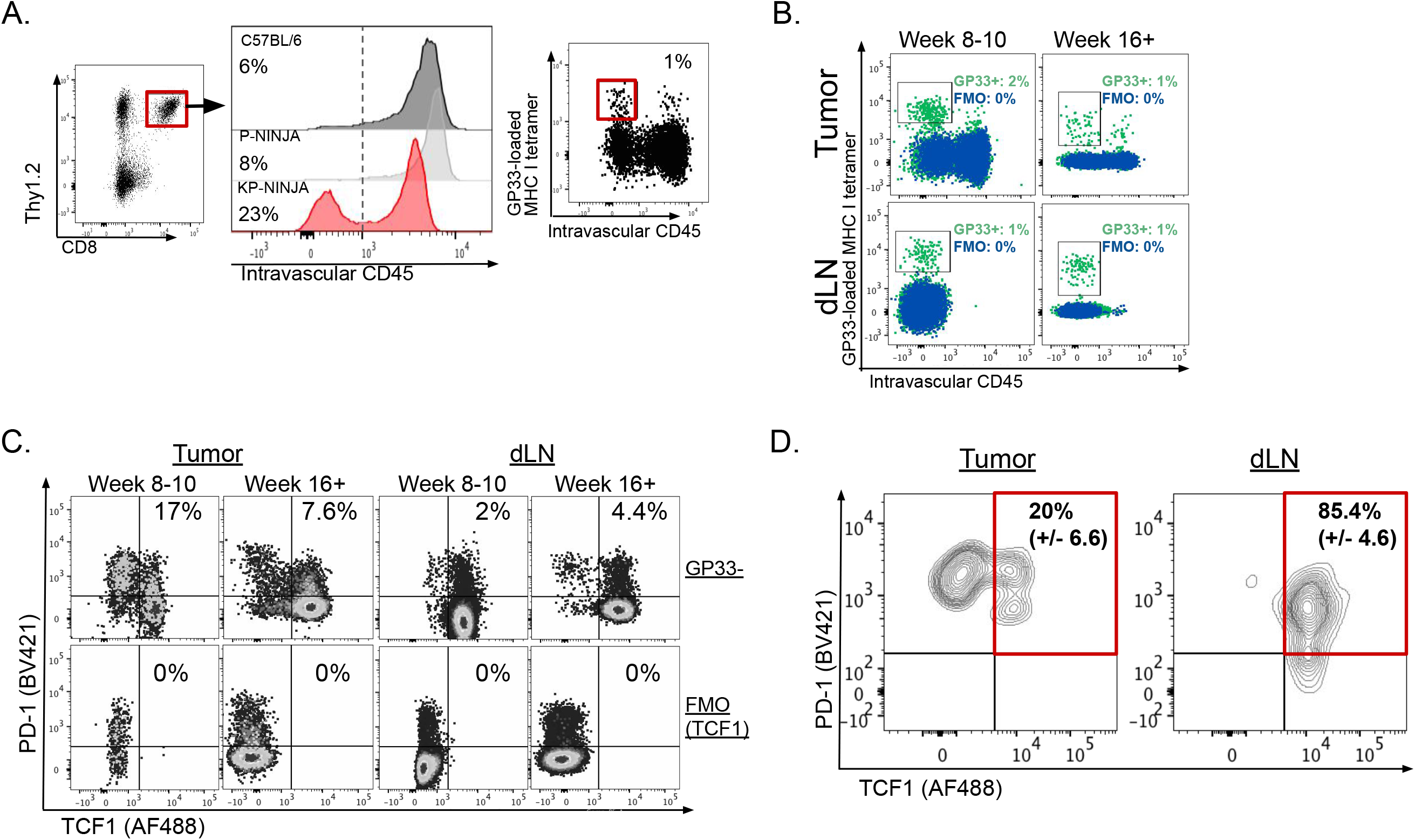
Representative plots from FACs. (A) KP-NINJA, P-NINJA, and C57BL/6 mice were infected intratracheally with 2.5 x 10^7^ PFU Ad5mSPC-Cre and treated with doxycycline and tamoxifen 7-10 days p.i. Lung tumors were harvested 8 weeks p.i. and intratumoral, tumor-specific CD8^+^ T cells were analyzed by flow cytometry. Dotted line reflects %GP33-specific CD8^+^ T cells in C57BL/6 control mice. (B) Representative flow cytometry dot plots demonstrating GP- 33-loaded MHC I tetramer staining and GP-33-specific gate (blue; pre-gated on singlets, THY1.2^+^ ivCD45-, CD8^+^) drawn based on FMO control (green) for early (left) and late (right) tumors (top) and dLN (bottom). (C) Representative flow cytometry dot plots displaying extracellular expression of PD-1 and intracellular expression of TCF1 on tissue GP33-negative CD8^+^ T cells (top) and FMO control for TCF1 (bottom) from the tumors (left) and dLNs (right) at early (8-10 weeks) and late (16+ weeks) time points after infection. Cells pre-gated on i.v. CD45-Thy1.2^+^CD8^+^ singlets. (D) <u>KP</u> mice were infected with mClover-GP33-80-Cre lentivirus. Tumor-draining LNs and tumor-bearing lungs were harvested at 8 weeks pi and single cell suspensions were obtained and stained using the same antibody panel as in Figure 1A. Representative flow cytometry dot plots displaying extracellular expression of PD-1 and intracellular expression of TCF1 on tissue GP33-specific CD8^+^ T cells from the tumor (left) and dLN (right). Averages +/− SEM reported for percent of PD-1^+^TCF1^+^ (20 +/− 6.6% and 85.4 +/− 4.6%) of total GP33-tetramer^+^ CD8 T cells. Cells pre-gated on i.v. CD45-Thy1.2^+^CD8^+^ singlets; n=11.

**Figure S2:**
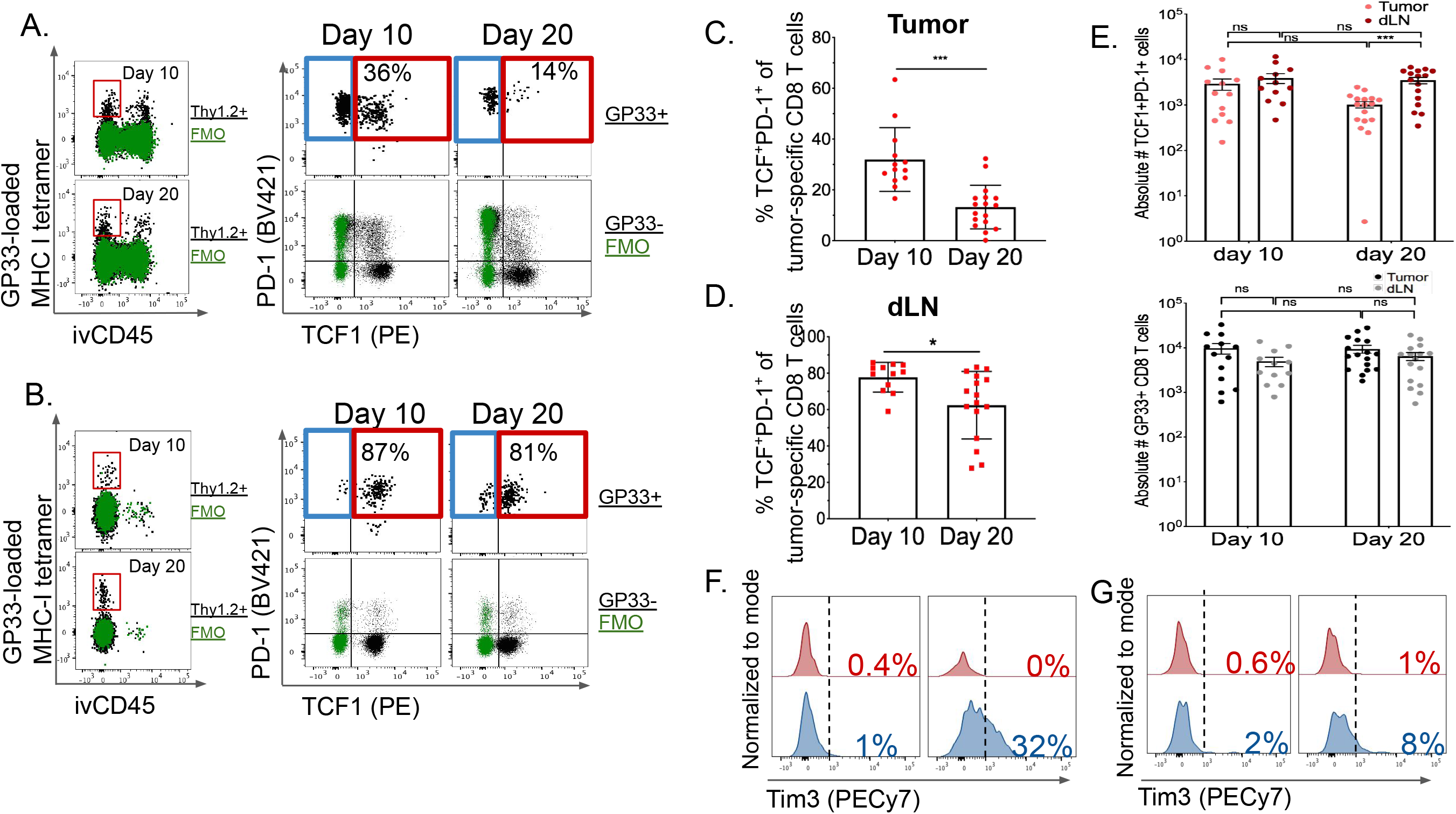
Tumor-specific TCF1^+^CD8^+^ T cells are present throughout disease progression in transplant KP-NINJA lung tumors. (A-F) 200k KPN1 cells were transplanted orthotopically via i.v. injection into adult C57BL/6 mice and tumor-specific CD8^+^ T cells were assessed 10 and 20 days post-injection in tumors (A) or tumor-dLN (B). (A-B) Representative dot plots showing GP-33- specific (GP-33-loaded MHC I tetramer^+^) gating based on FMO controls (green) at day 10 (top) and day 20 (bottom). Extracellular PD-1 and intracellular TCF1 staining of intratumoral tumor-specific CD8^+^ T cells, in the tumor (A) and dLN (B), pre-gated on singlets, Thy1.2^+^CD8^+^ i.v.CD45, and GP33-loaded MHC I tetramer^+^. PD-1 and TCF1 staining of GP33- CD8 T cells (black) and TCF-FMO controls (green). (C-D) Percent TCF1^+^PD-1^+^ T_SL_ of total tumor-specific CD8^+^ T cells in tumor (C) and dLN (D) from day 10 to 20 p.t. (C, p=0.0062; D, p=0.0047). (E) Absolute numbers of TCF1^+^PD-1^+^ T_SL_ (top; ***p=0.0006) and total GP-33-specific CD8 T cells (bottom) in tumors (light red and black, respectively) and dLNs (dark red and gray, respectively). Data from 5 independent experiments: n=13 tumors at 10 days post-injection;n=17 tumors at 20 days post injection; n=12 dLN at 10 days post injection;n=16 dLN at 20 days post-injection. Statistics based on two-tailed, unpaired t-test. (F-G) Representative histograms displaying extracellular Tim3 expression of TCF1^-^PD-1^+^ vs. TCF1^+^PD-1^+^ tumor-specific CD8^+^ T cells at early (10 days) and late (20 days) after tumor injection in tumors (F) and dLNs (G).

**Figure S3:**
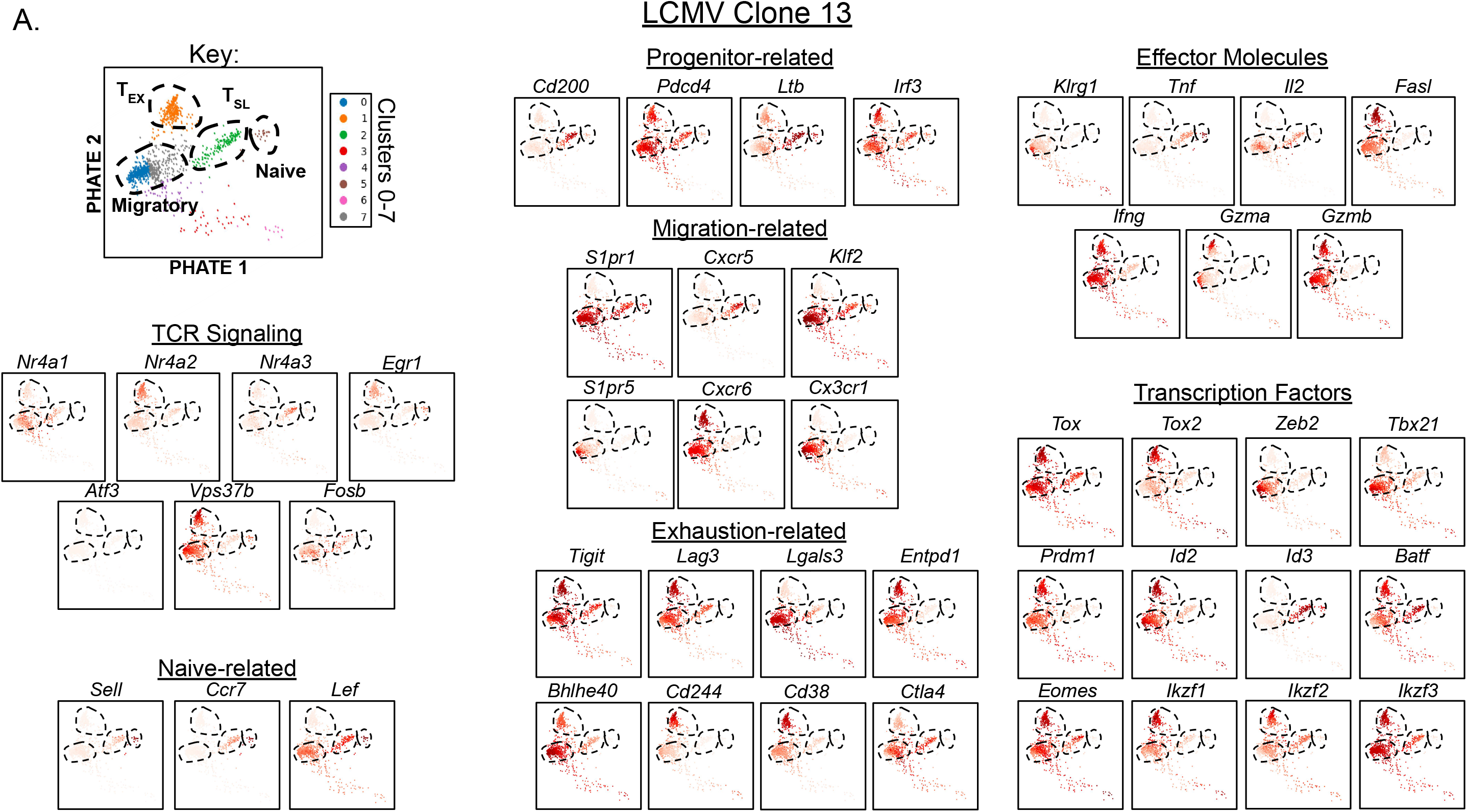
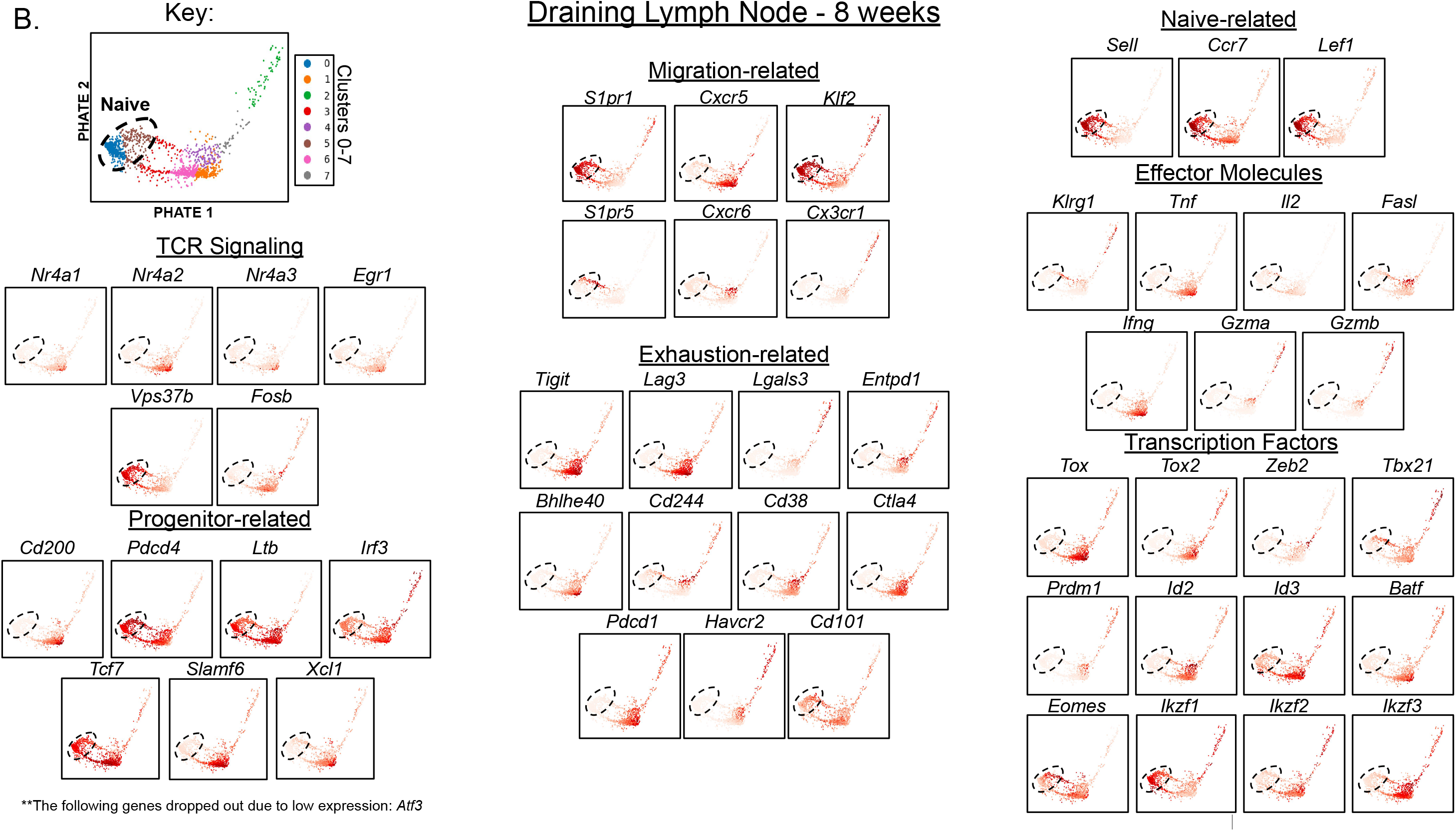
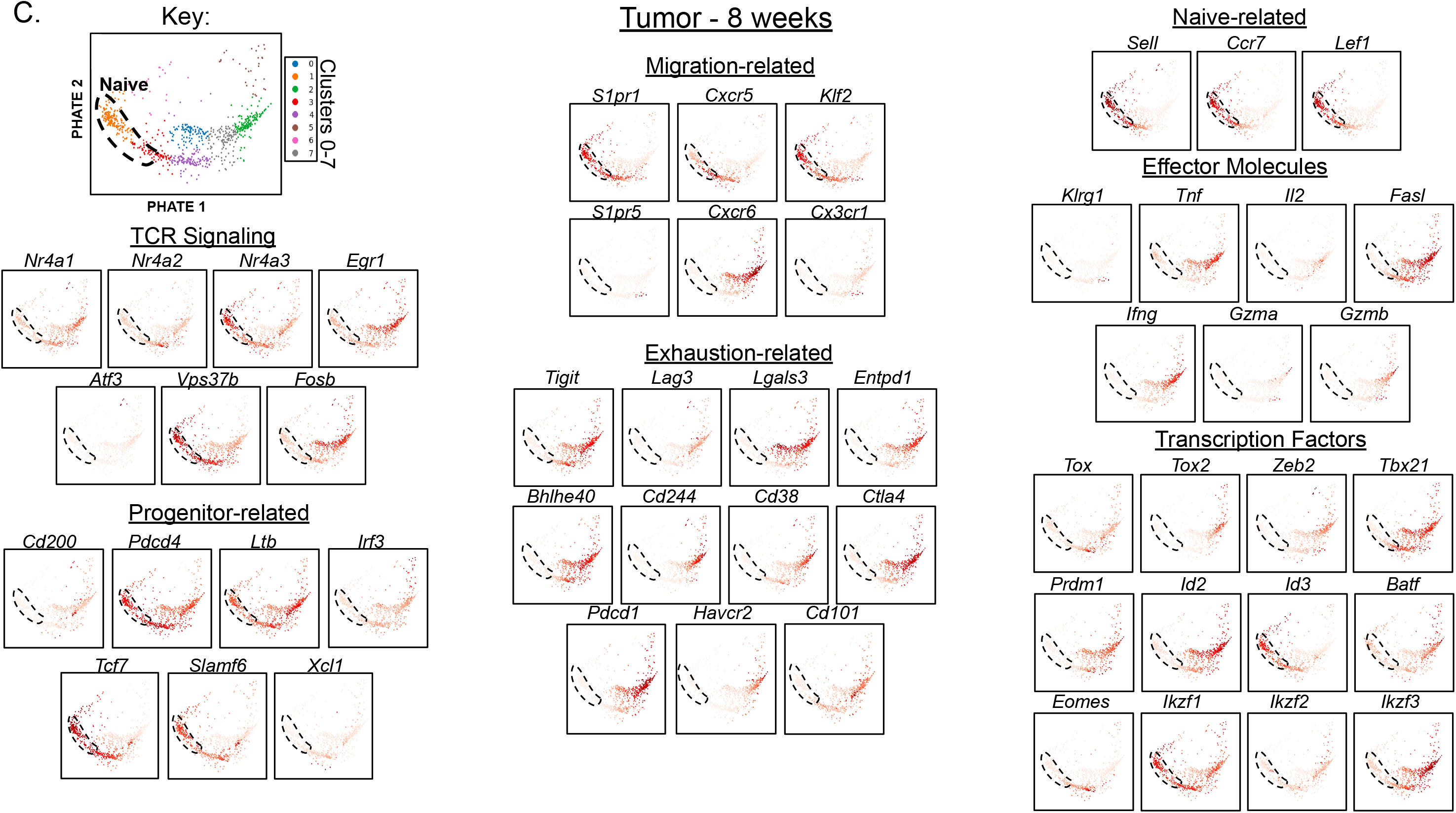
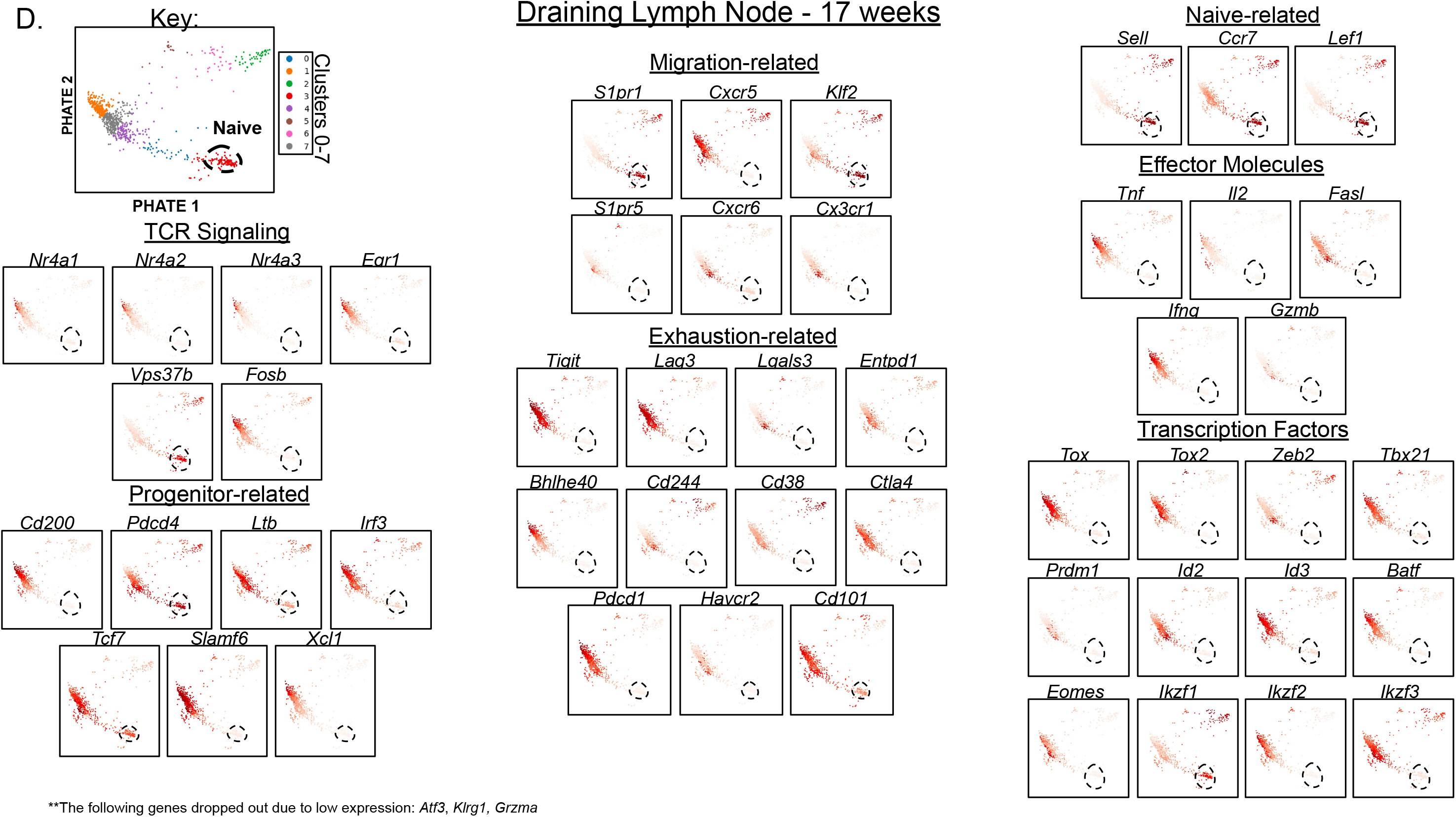
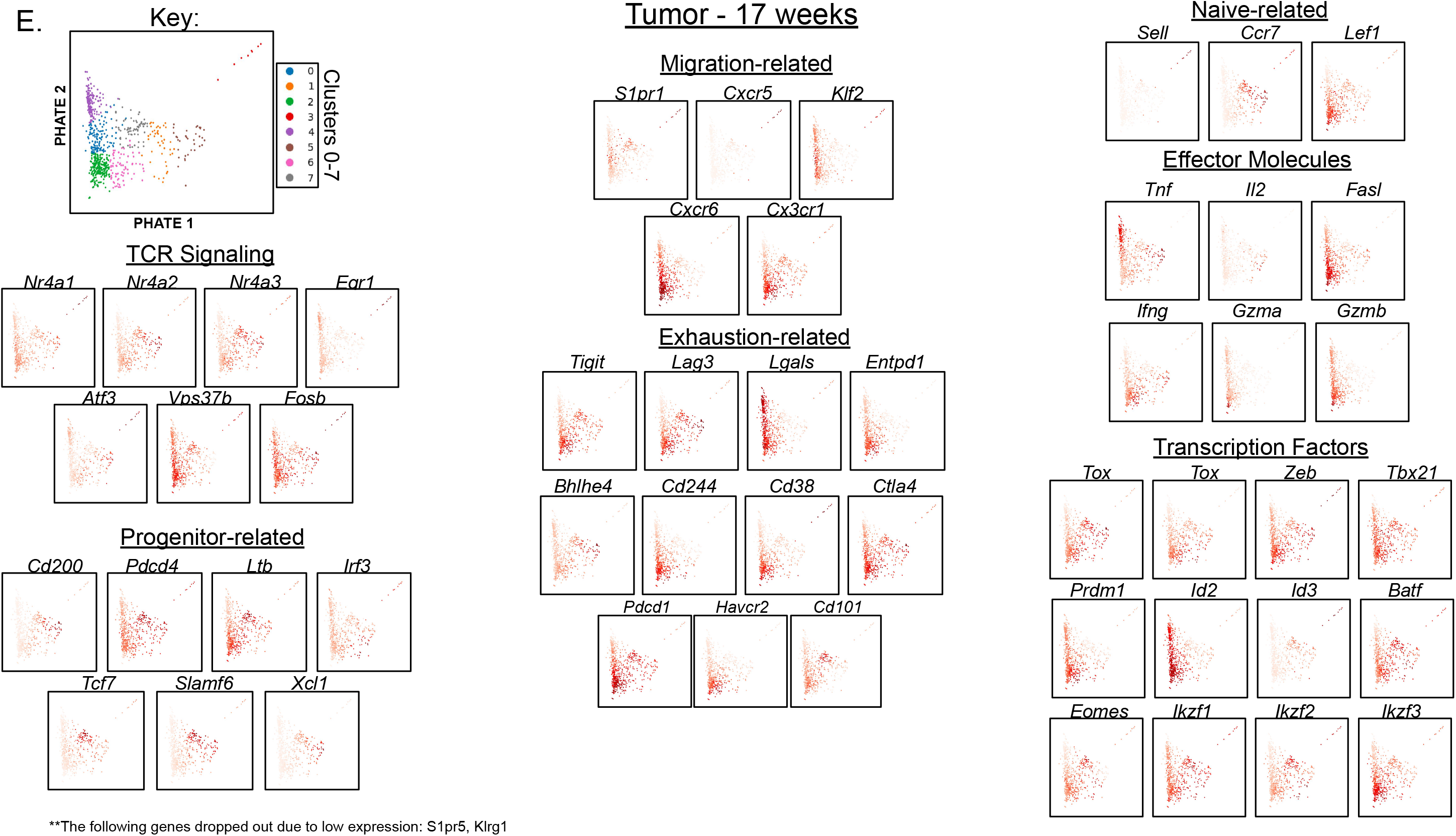
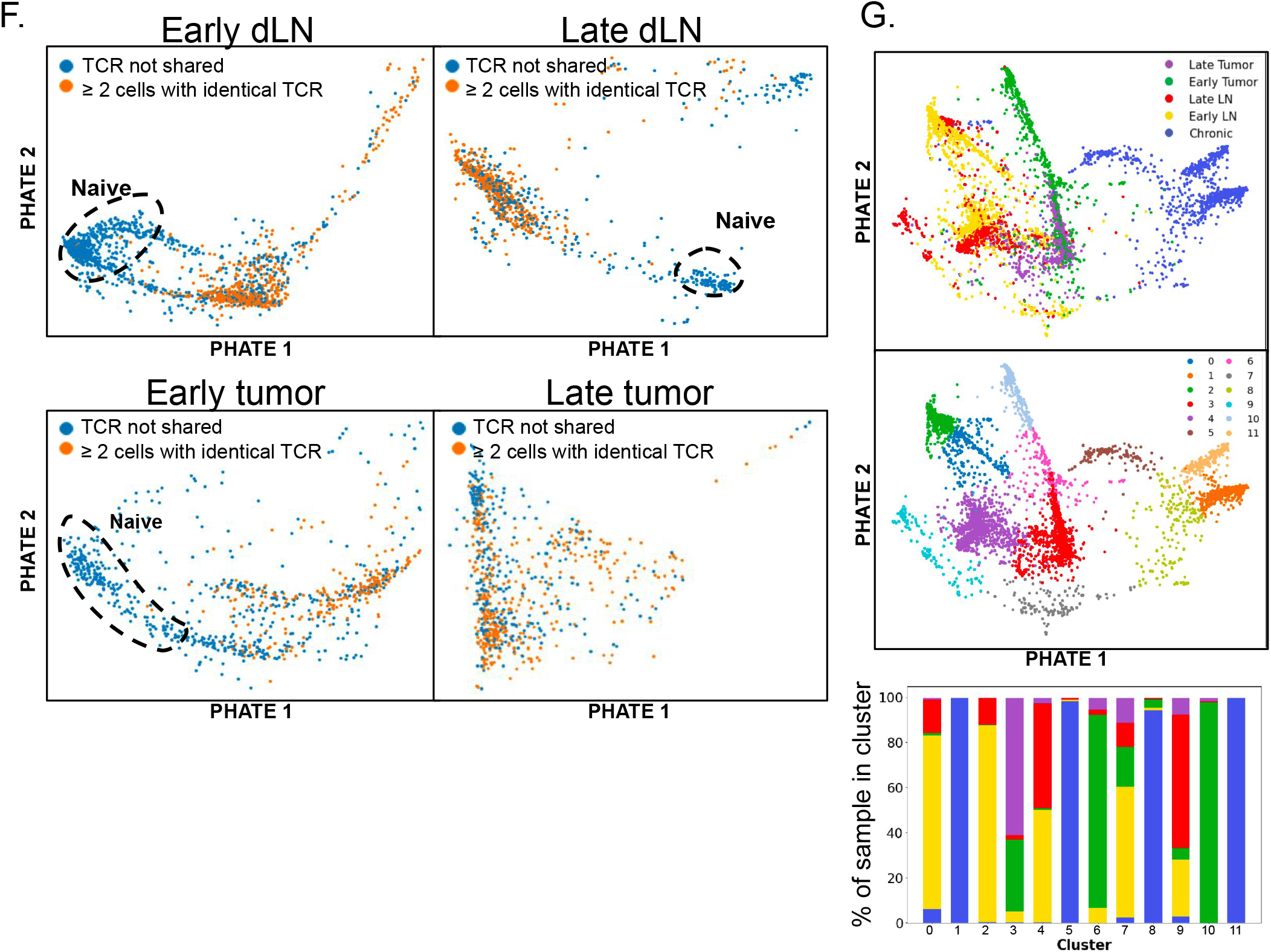
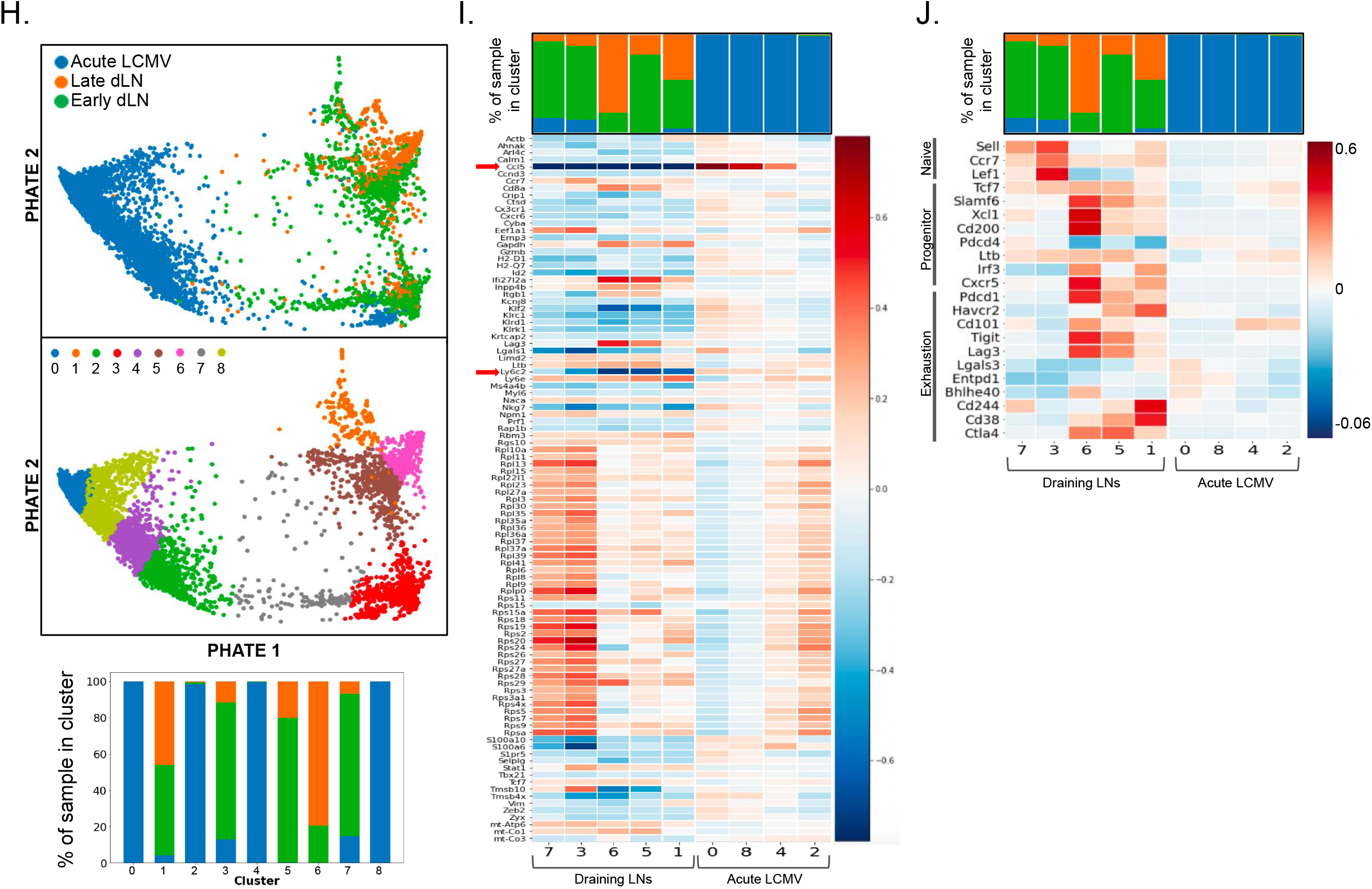
Gene expression plots for LCMV, early and late dLNs and tumor comparisons. (A-J) GP33-specific CD8^+^ T cells were sorted from various tissues (i.v.CD45^-^ CD8^+^GP33-loaded MHC I tetramer^+^) and submitted to the Yale Center for Genome Analysis for 10x single cell RNA-sequencing. Spleens were harvested from C57BL/6 mice 28 days following infection with LCMV-Clone 13 or LCMV-Armstrong. Tumors and dLNs were harvested from tumor-bearing KP-NINJA mice 8 and 17 weeks after intratracheal infection with Ad5mSPC-Cre and doxycycline and tamoxifen treatment. Gene expression profiles for GP33-specific CD8^+^ T cells from chronic LCMV-clone 13 infection (A), early (8 weeks p.i.) dLN (B), early tumor (C), late (17 weeks p.i.) dLN (D), and late tumor (E) visualized using PHATE maps. Expression profiles of additional progenitor-related (*Cd200, Pdcd4, Ltb, Irf3, Tcf7, Slamf6, Xcl1*), migration-related (*S1pr1, Cxcr5, Klf2, S1pr5, Cxcr6, Cx3cr1)* and terminally exhausted (*Tigit, Lag3, Lgals, Entpd1, Bhlhe4, Cd244, Cd38, Ctla4, Pdcd1, Havcr2, Cd101*) genes, as well as key effector cell-associated (*Klrg1, Tnf, Il2, Fasl, Ifng, Gzma, Gzmb*) and transcription factor (*Tox, Tox2, Zeb2, Tbx21, Prdm1, Id2, Id3, Batf, Eomes, Ikzf1, Ikzf2, Ikzf3*) genes, genes associated with intracellular TCR signaling (*Nr4a1, Nr4a2, Nr4a3, Egr1, Atf3, Vps37b, Fosb*) and a naïve cell phenotype (*Sell, Ccr7, Lef1; right top*). **In some samples, expression profiles for certain genes have been omitted due to drop out caused by low expression levels. For LCMV-Clone 13 (A), please refer to Figure 2C for additional expression plots. (F) PHATE map visualizing TCR abundance based on single cell TCR-sequencing for early (left) and late (right) dLNs (top) and tumors (bottom). (G) 2- dimensional PHATE map of GP33-specific CD8 T cells from chronic LCMV infection, early dLN, late dLN, early tumor, and late tumor combined (top) and divided into 12 (0-11) clusters (bottom) with distribution of samples for each clusters shown below. (H-J) Single cell-sequencing results of GP33-specific CD8^+^ T cells from acute LCMV (Armstrong) infection (10,768 cells analyzed; 12,960 genes detected), and from early and late dLNs were co-embedded. (H) Cells were visualized by PHATE map (top) and clustered 0-8 (bottom),with distribution of clusters between samples shown below. (I) Heat map displaying top differentially expressed genes between acute LCMV and early dLN visualized for each cluster based on analysis from H. (J) Heat map displaying gene expression for Naïve-, progenitor-, and exhaustion-related genes.

**Figure S4:**
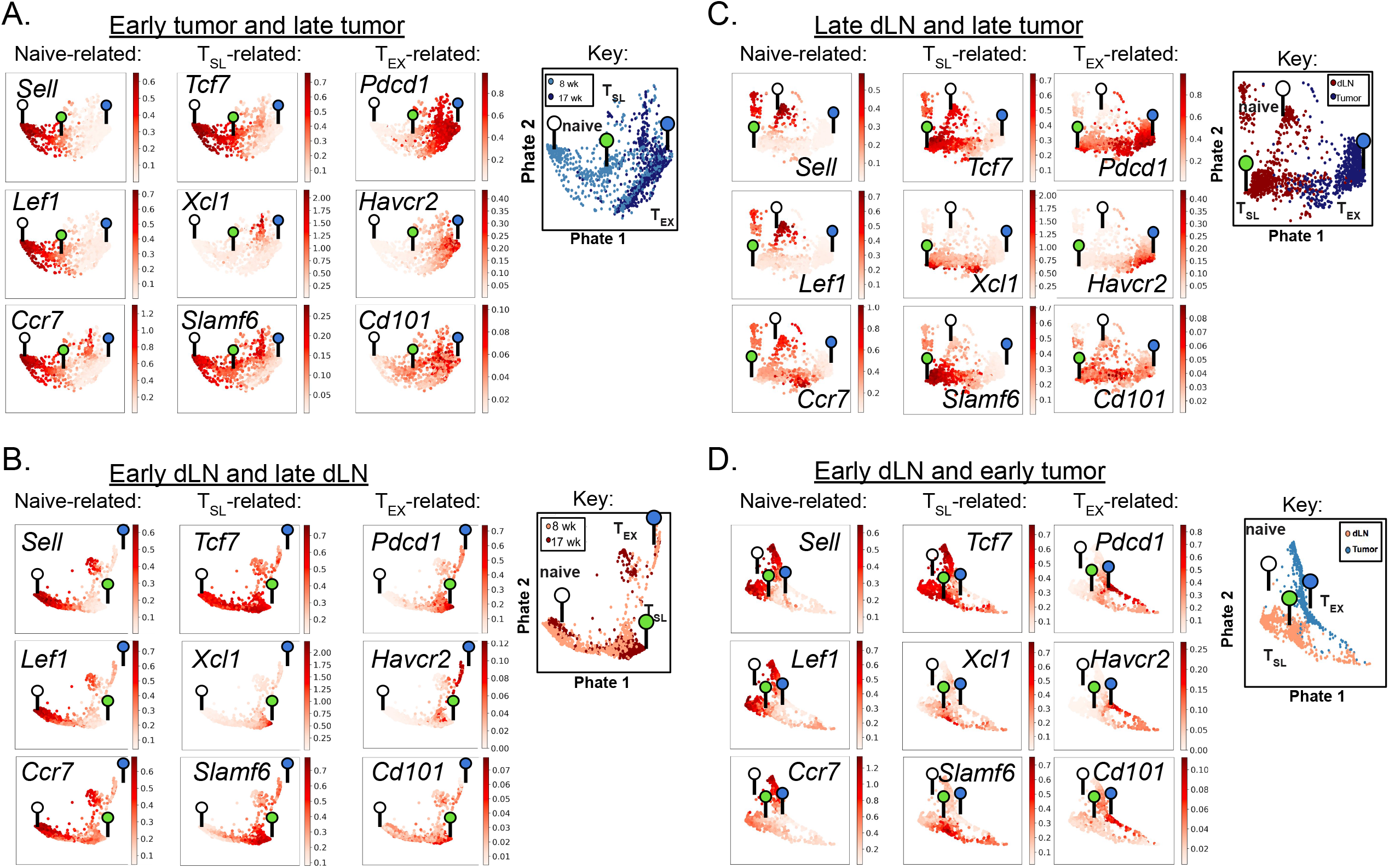
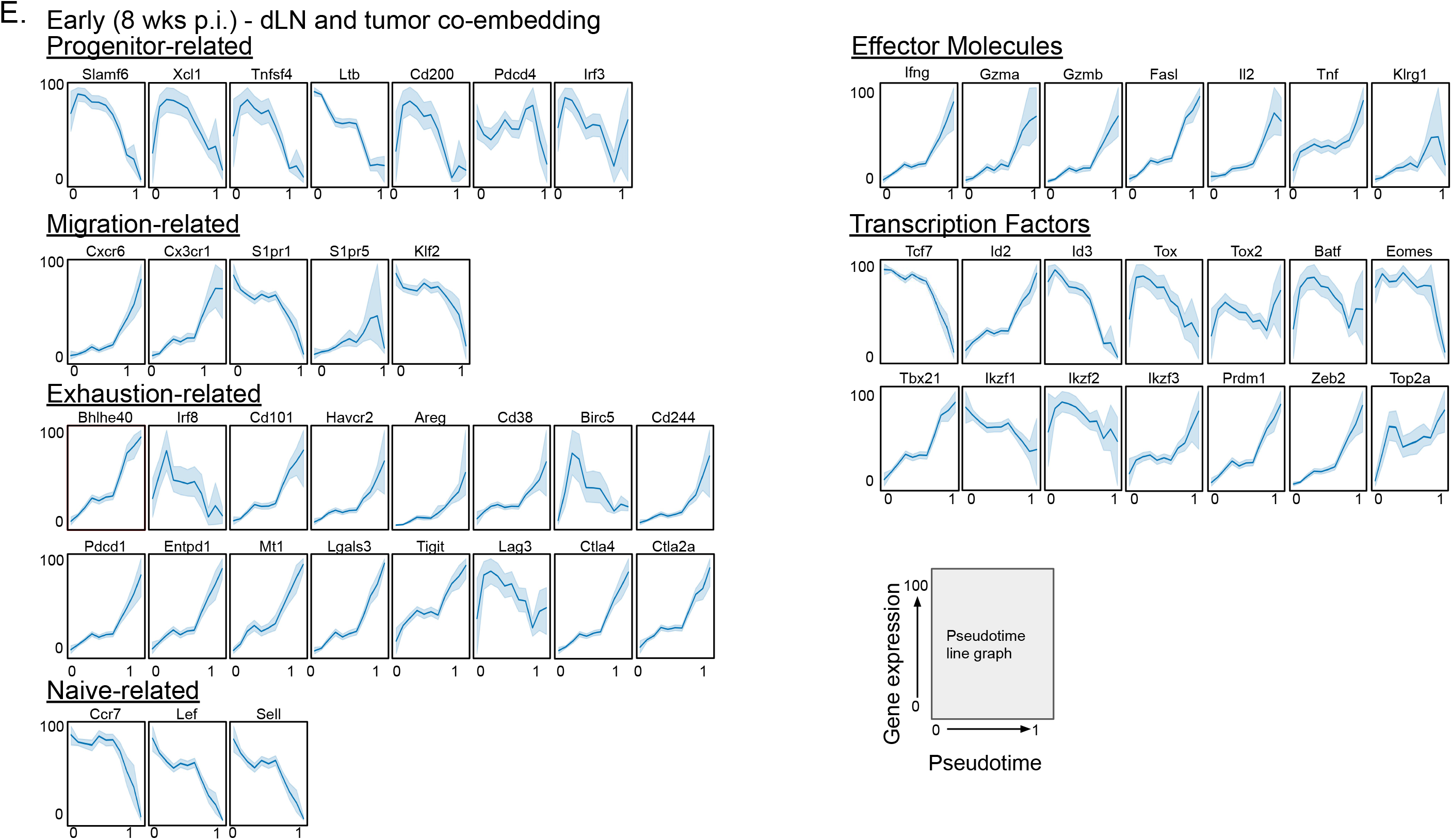
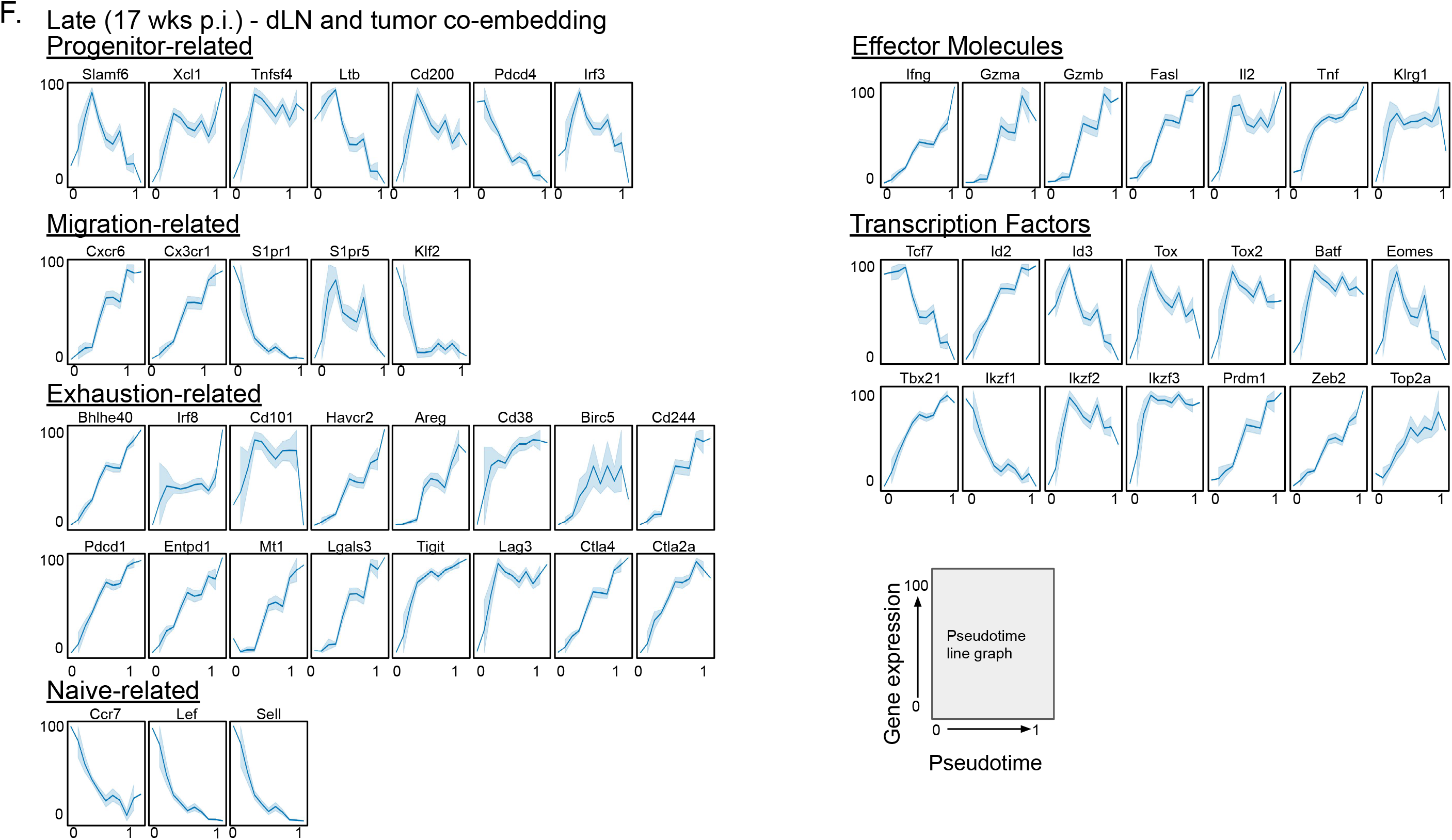
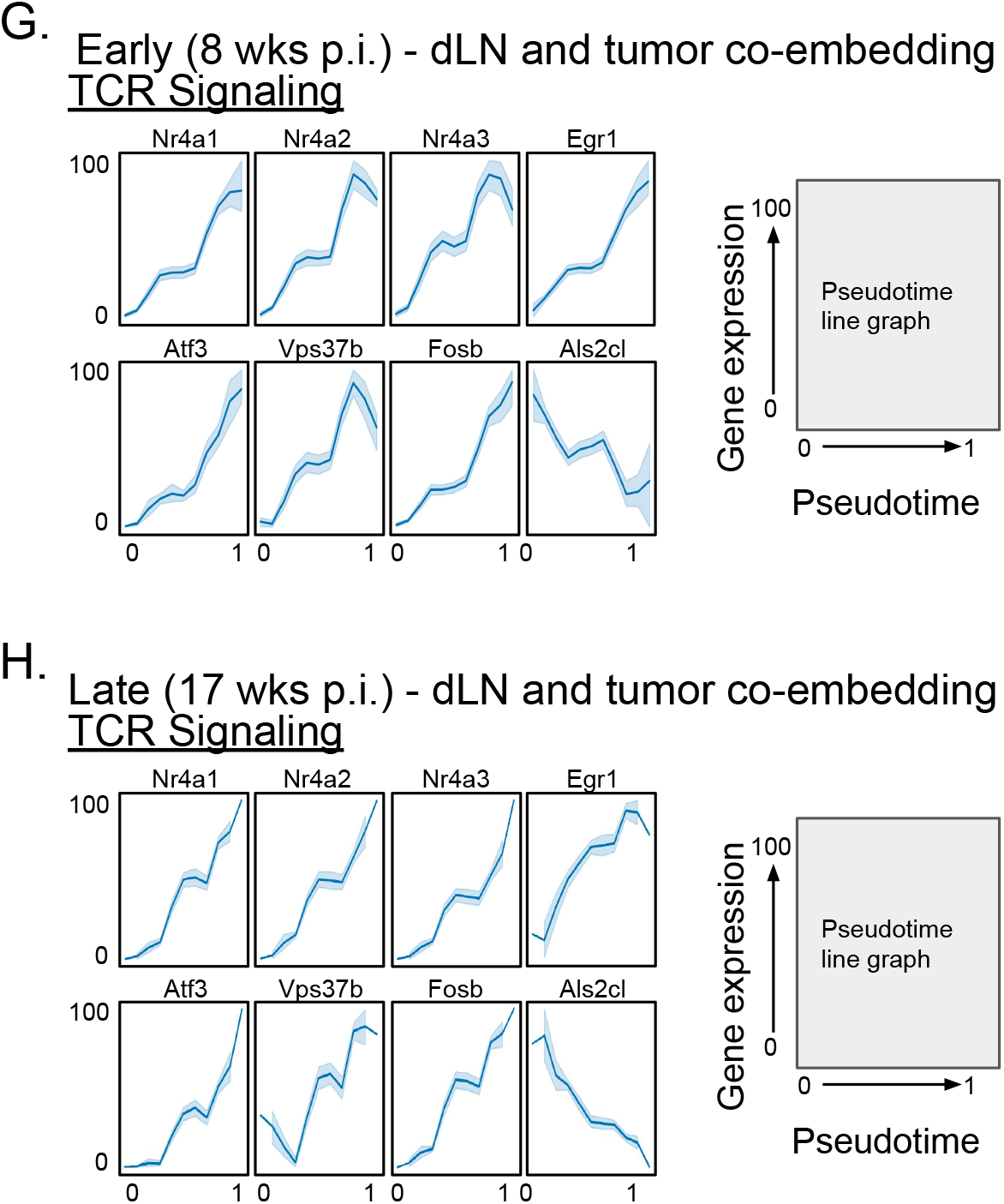
Gene expression and transcript dynamics for co-embedded samples. (A-D) Single-cell transcriptomics of tumor-specific CD8^+^ T cells from early (8 weeks p.i.) tumors and late (17 weeks p.i.) tumors (A), early dLNs and late dLNs (B), late dLNs and late tumors (C), and early dLNs and early tumors (D) from KP-NINJA mice were co-embedded and expression profiles were visualized by PHATE maps for key naïve-related (*Sell, Lef1, Ccr7*; left), T_SL_-related (*Tcf7, Xcl1, Slamf6*; center), and T_EX_-related (*Pdcd1, Havcr2, Cd101*; right) genes. The location of transcriptional signatures for the major cell states identified (Naïve (white), T_SL_ (green), and T_EX_ (blue)) are indicated by markers on pseudotime visualizations. (E-H) Pseudotime line graphs for tumor-specific CD8^+^ T cell co-embedding for early (8 weeks p.i.) dLN and early tumor (E, G) and for late (17 weeks p.i.) dLN and late tumor (F, H), showing progenitor-related (*Slamf6, Xcl1, Tnfsf4, Ltb, Cd200, Pdcd4, rf3*), migration-related (*Cxcr6, Cx3cr1, S1pr1, S1pr5, Klf2*), exhaustion-related (*Bhlhe40, Irf8, Cd101, Havcr2, Areg, Cd38, Birc5, Cd244, Pdcd1, Entpd1, Mt1, Lgals3, Tigit, Lag2, Ctla4, Ctla2a*), naïve-related (*Ccr7, Lef1, Sell*) genes, as well as effector cell-related (*Ifng, Gzma, Gzmb, Fasl, Il2, Tnf, Klrg1*), transcription factor (*Tcf7, Id2, Id3, Tox, Tox2, Batf, Eomes, Tbx21, Ikzf1, Ikzf2, Ikzf3, Prdm1, Zeb2, Top2a*) genes (E-F), and genes associated with TCR-signaling (*Nr4a1, Nr4a2, Nr4a3, Egr1, Atf3, Vps37b, Fosb, Als2cl*) (G-H). Velocity confidence estimates by scVelo were calculated for pseudotime analyses and were greater than 50 in all samples.

